# Unraveling Structural Disparities in Human and Mycobacterium Tuberculosis Type-I Fatty Acid Synthase

**DOI:** 10.1101/2024.07.17.603935

**Authors:** Akhil Kumar, Priyanka Rama, Harshwardhan H. Katkar

## Abstract

Type-I Fatty acid synthase is an essential enzyme present in diverse species including humans (hFAS-I) and mycobacterium tuberculosis (MtbFAS-I), and is an attractive antibacterial drug target. A structural comparison of the two enzymes is essential in order to identify selective drug targets in MtbFAS-I. In this work, we have analyze equilibrium average structures of hFAS-I and MtbFAS-I obtained from 100 ns long molecular dynamics simulation trajectories. Our structural analysis revealed that two of the seven domains present in both hFAS-I and MtbFAS-I, *viz*. dehydratase (DH) and enoyl reductase (ER), are significantly dissimilar. We further compared corresponding catalytic pockets in these two domains and analyzed their physicochemical characteristics. In addition to being large in MtbFAS-I, the pockets are significantly different in their physicochemical characteristics and water content.

## 1 INTRODUCTION

Type-I Fatty Acid Synthase (FAS-I) is a multi-domain complex that plays a crucial role in *de-novo* synthesis of fatty acids, which are a major component of the cellular membrane [1, 2, 3]. The presence of FAS-I in bacteria, fungi and mammals underscores its significance across diverse species, where it actively supports essential cellular processes [4, 5]. FAS-I adopts a variety of structures across different species. Mammalian FAS-I (mFAS-I) is a homodimer with an open X-shaped structure. All of its functional domains reside in two identical 2500-amino-acids-long polypeptide chains. The structure of FAS-I from fungal species *Thermomyces lanuginosus* [6], *Saccharomyces cerevisiae* [7] (SC), *Candida albicans* [8] and bacterial species *Mycobacterium smegmatis* [9] and *Mycobacterium tuberculosis* [10] (Mtb) is homohexameric, with a closed-barrel shape. Seven functional domains reside on each of six identical 3092-amino-acids-long polypeptide chains in Mtb. In fungal FAS-I, these domains are arranged in two separate polypeptide chains, *viz. α* and *β*, with 1878 and 2060 amino acid residues, respectively. The mFAS-I lacks a rigid scaffold structure with loosely connected domains, imparting significant structural flexibility to the complex [11]. About 9% of its sequence is dedicated to exposed linkers [11]. About 35% of MtbFAS-I and 50% of SC FAS-I sequence is involved in scaffolding of their respective structure [10]. Organization of domains along the primary sequence is different in MtbFAS-I and mFAS-I [11, 12]. Despite these variations in the structure of FAS-I across different species, essential features of the fatty acid synthesis reaction mechanism are conserved and are described below. [13].

Synthesis of fatty acid involves three stages: initiation, elongation and termination. The initiation stage involves condensation of acetyl and malonyl moieties at the Ketoacyl synthase (KS) domain. During the elongation stage, four successive reactions take place multiple times (seven times to produce palmatic acid in Mtb), until the growing fatty acid reaches the desired length. Each elongation cycle involves addition of 2 carbon atoms provided via a malonyl moiety that are added to the growing substrate. The substrate is sequentally shuttled between four enzymatic domains, KS, ketoacyl reductase (KR), dehydratase (DH) and enoyl reductase (ER) [14, 15, 16, 17, 18]. Termination occurs once the desired length of carbon atoms are added to the growing fatty acid substrate.

In MtbFAS-I, two distinct domains are involved in the initiation and termination stages. Acetyltransferase (AT) transfers acetyl moiety to KS during the initiation stage. Malonyl/palmitoyl transferase (MPT) transfers the malonyl moiety during the initiation and elongation stages, and also terminates the reaction. Conversely, in mFAS-I, malonyl/acetyl transferase (MAT) is responsible for transferring both, acetyl and malonyl moieties to KS [19, 20, 21]. A distinct thioesterase (TE) domain terminates the reaction to release the synthesized palmitic acid [13, 22].

The shuttling of the growing substrate between multiple domains is facilitated by the mobile Acyl Carrier Protein (ACP) domain that is connected to the rest of the complex through two linkers [23, 24].

All mammalian FAS-I ACPs, including hFAS-I ACP, consist of four *α* −helices, while MtbFAS-I ACP consists of a total of eight *α* −helices: four forming the catalytic core and four forming the structural core [25, 26, 27]. In MtbFAS-I, two linkers, *viz*. the peripheral linker (PL) connecting MPT to ACP, and the central linker (CL) connecting ACP to KR, composed of about 51 and 21 residues respectively, play a crucial role in the shuttling process [28, 29]. Similarly, in hFAS-I, ACP is connected via two linkers. A longer linker connecting ACP and TE comprises about 25 residues, and a shorter second linker connecting ACP and KR is composed of about 17 residues [30]. Although MtbFAS-I and hFAS-I have ACPs that are connected via two linkers, ACP is more mobile in hFAS-I as its connected via the longer linker with the mobile TE domain. In contrast, ACP’s motion is constrained by double-tethering in MtbFAS-I [11]. In Mtb, PL has high Proline and Alanine content in comparison to the corresponding linker in hFAS, resulting into a stiffer linker [11]. Nonetheless, owing to their high flexibility relative to stable domains in the complex, linkers have not been resolved in the latest high-resolution structures of the entire hFAS-I and MtbFAS-I complexes [12, 13]. Changes in the length of hFAS-I’s ACP-TE linker have been shown to affect the catalytic activity marginally, with about 30% reduction in activity of the complex upon complete deletion of the linker [31]. A similar reduction in the probability of accessing domains has been shown in fungal FAS-I using coarse-grained simulation [28]. A unique set of PL conformations were reported in molecular dynamics simulation of ACP persistently interacting with DH in MtbFAS-I complex [29].

In earlier reports, the mammalian FAS-I was discovered to be a dimer composed of two polypeptide chains positioned adjacent to one another [32, 33]. In 2002, the first low-resolution three dimensional structure of hFAS-I was resolved using cryo-electron microscopy (cryo-EM) [34]. However, the orientation of the two chains in the complex was not conclusively assigned in this structure, as models with and without imposed symmetry were found to converge with the limited resolution data. Although initial models assumed a head-to-tail structural arrangement of domains between the two chains, this was improved in subsequent studies to establish that a head-to-head structural arrangement of the two chains exists [35]. The first crystal structure of porcine FAS-I complex was resolved at a near-atomic resolution of 4.5 Å, providing direct evidence of the head-to-head structural arrangement of the two chains [36]. An improved resolution structure of porcine FAS-I at 3.2 Å revealed the presence of two additional non-enzymatic domains [11]. Due to the considerable mobility of TE and ACP, both of these otherwise complete structures of the FAS-I complex had missing TE and ACP domains, and unstructured linkers. The structure of isolated TE domain of hFAS-I was resolved at 2.6 Å and later at 2.3 Å using X-ray diffraction [37, 38]. Also, the structure of hFAS-I ACP in a complex was resolved at 2.7 Å using X-ray diffraction [25]. A homology model of hFAS-I was generated in Ref. [30] using the 3.2 Å resolution structure of porcine FAS-I [11] and docking the available high resolution structures of hFAS-I TE and hFAS-I ACP domains [38, 25], with additional refinement. Computational docking are used to characterize the ligand-binding pocket of hFAS-I TE domain ad their inhibitors. [39].

Among fungal FAS-I, the structure of *Thermomyces lanuginosus* FAS-I complex was initially reported at 5 Å and further refined at 3.1 Å using X-ray diffraction, with all domains resolved except the ACP and phosphopantetheinyl transferase (PT) domains [40]. The structure of SC FAS-I, reported at 4 Å using X-ray diffraction, was inclusive of ACP stalled at KS inside the closed complex, and PT on the exterior surface of the complex [7]. An improved structure of SC FAS-I with ACP stalled at the KS domain was recently reported at a resolution of 1.9 Å using cryo-EM, with the additional structure of ACP stalled at KR, ER, MPT, and AT domains determined at coarser resolutions [41].

The structure of bacterial FAS-I was first reported for *Mycobacterium Smegmatis* using cryo-EM data with imposed symmetry at a resolution of 7.5 Å. The structure revealed a significant similarity with the yeast FAS-I structure, with lack of a PT domain. Further analysis of *Mycobacterium Smegmatis* FAS-I from Ref. [9] and cryo-EM data of MtbFAS-I without any imposed symmetry at 20 Å revealed that compared to the rigid fungal FAS-I complex with large number of scaffolding residue contacts, bacterial FAS-I exhibits a more flexible structure [10]. A detailed, near-atomic resolution structure of MtbFAS-I was recently reported at 3.3 Å using cryo-EM [12]. ACP and linkers were not resolved in this structure owing to the mobility of ACP and flexibility of linkers. However, electron clouds were observed near KS, albeit at a lower resolution. In our effort to study the MtbFAS-I dynamics, our laboratory has previously reported a homology model of MtbFAS-I complex that includes ACP and linkers. Our model was generated using the cryo-EM MtbFAS-I structure at 3.3 Å from Ref.[12] and the 3.1 Å resolution cryo-EM structure of SC FAS-I [42] with ACP stalled at KS as template for homology modeling, followed by loop modeling of the linkers. The model was further refined using the 5.1 Å resolution cryo-EM structure of ACP bound to DH [43] for obtaining a structure of MtbFAS-I with ACP stalled at DH.

With the availability of accurate models for complete hFAS-I and MtbFAS-I complex a detailed comparison of their structure and dynamics is now possible. Such a comparison can provide important insights that can be crucial for the rational design of drugs and for identifying novel targets in MtbFAS-I.

In this work, we analyze the structure and dynamics of the hFAS-I and MtbFAS-I complexes with the goal of gaining detailed structural insights. We performed molecular dynamics simulation of the two complexes using the homology models of hFAS-I and MtbFAS-I from Refs. [30] and [29] respectively. In Section 3, we report a superimposition of the average structures of functionally-similar domains from the two trajectories, with a focus on relative positions and orientations of catalytic residues. Dynamics of corresponding residues in these domains are also presented. We identify DH and ER as structurally distinct domains and report key characteristics of their catalytic pockets. Our findings warrant a detailed investigation of the possibility of using DH and ER as selective therapeutic targets.

## 2 METHODS

### 2.1 Molecular Dynamics Simulation

Coordinates of homology model of hFAS-I complex were retrieved from Ref. [30]. GROMACS (RRID:SCR_014565) version 2020.6 was used to prepare the system and to perform molecular dynamics simulation, with CHARMM36 force-field. [44, 45]. The retrieved homology model was solvated using TIP3 [46] water model and a 370Å×185Å×209Å simulation box. The solvated system was neutralized by adding 76 Na^+^ ions. The final system had 1416663 atoms, consisting of 76592 protein atoms, 1339995 water atoms and 76 Na^+^ ions. To reduce overlapping of atoms in the system, energy of the prepared system was minimized using the steepest-descent algorithm with a force tolerance of 10000 kJ/mol^−1^ nm^−1^. The resulting system was gradually heated from 0 K to 310 K in 3 ns with a linearly increasing temperature, followed by 2 ns of equilibration at a constant volume and temperature using the NVT ensemble with a V-rescale thermostat and harmonic restraints on positions of all hFAS-I atoms with a force constant of 1000 kJ mol^−1^ nm^−2^. Subsequently, 5 ns of equilibration at a constant pressure of 1 bar and temperature of 310 K was performed using a V-rescale thermostat and a Parrinello-Rahman barostat, with harmonic restraints on backbone atoms of hFASI with a force constant of 1000 kJ mol^−1^ nm^−2^. Following this, a production run was performed to generate a 100 ns-long trajectory of the hFAS-I complex without using any position restraints. An average structure of the complex was calculated using this 100 ns-long trajectory.

A 100 ns simulation trajectory of the relatively large MtbFAS-I from our previous work was utilized in this study with details available in Ref. [29]. Briefly, starting from the cryo-EM structure of MtbFAS-I with missing PL, CL and ACP (PDB ID 6GJC) [12], a model of MtbFAS-I was generated using the structure of a homologous ACP from SC (PDB ID 6TA1) [7] and loop modelling [47]. The model was solvated using a box of 300Å × 300Å × 300Å and TIP3P water, ionized, energy minimized, and gradually heated, using a similar protocol as described above for hFAS-I. Equilibration was performed in two stages: 5 ns of NVT equilibraition followed by 21 ns of constrained equilibration at a pressure of 1 bar and temperature of 310 K with Parrinello-Rahman barostat and V-rescale thermostat (NPT ensemble), with slowly relaxing position restraints on various MtbFAS-I atoms. Target MD simulation was performed twice in order to move ACP near DH; the first simulation based a target structure of ACP near DH (PDB ID 6WC7), and the refined second simulation using a unique PL conformation explored in the first target MD simulation. A production run was performed to generate the 100 ns-long trajectory of the MtbFAS-I complex. Average structure of the complex was calculated using this 100 ns-long trajectory.

A six-chain average structure of MtbFAS-I and a two-chain average structure of hFAS-I were generated by calculating time-averaged and chain-averaged coordinates of the protein complex from the corresponding 100 ns long trajectory. Each chain in each frame was aligned to chain A in the first frame of the trajectory before calculating the average coordinates.

### 2.2 Structure and Sequence Comparison

The MultiSeq tool in VMD (RRID:SCR_024368) utilizing STAMP algorithm was used for aligning average structures of hFAS-I and MtbFAS-I, based on three dimensional structural fitting of corresponding residues in the two sequences [48, 49]. A cutoff of 0.5 was used for the Q_*res*_ value. Q_res_ represents the individual contribution of each residue to the total Q_*H*_ value of aligned structures; the latter being a measure of overall structural similarity

### 2.3 Pocket identification

The average domain structures were used to predict pockets using the CASTp3 web server [50]. Of the multiple pockets predicted for each domain, the pocket with largest volume that included all of the domain’s catalytic residues listed in Table 1 was selected as the catalytic pocket. All catalytic-pocket forming residues were used to calculate the volume of catalytic pocket using the tool trj_cavity [51]. The pocket volume of DH and ER presented in sections **??** and **??** are total volume calculated with trj_cavity for all the pocket forming residue predicted via CASTp3 server. Identified pockets were visualized using UCSF ChimeraX (RRID:SCR_015872).

**TABLE 1.** Catalytic residues present in each domain in hFAS-1 and MtbFAS-I with their alignment score, RMSD, Percentage Identity.

| Domain<br>hFAS-I/MtbFAS-I | Catalytic residues<br>hFAS-I | Catalytic residues<br>MtbFAS-I[6] | $Q_H^+$<br>0-1 | RMSD<br>Å% |
| --- | --- | --- | --- | --- |
| MAT/AT<br>(415-820/1-366)<br>(111%) | SER581 [52]<br>HIS683 | SER128<br>HIS266 | 0.48 | 2.77 |
| MAT/MPT<br>(415-820/1311-1709)<br>(102%) | HIS683[53]<br>SER581 | HIS1545<br>SER1417 | 0.47 | 2.57 |
| KS/KS<br>(1-460/2388-3015)<br>(73%) | CYS161 [54, 55, 56, 57]<br>HIS331<br>HIS293 | CYS2699<br>HIS2851<br>HIS2892 | 0.46 | 2.43 |
| ACP/ACP<br>(2127-2193**/1757-1924)<br>(40%) | SER2159[25, 30] | SER1787 | 0.45 | 3.37 |
| KR/KR<br>(1879-2108*/2099-2387)<br>(80%) | SER2021 [58, 59]<br>TYR2034<br>LYS1995 | SER2253<br>TYR2265<br>LYS2269 | 0.45 | 3.47 |
| DH/DH<br>(858-1103/1028-1310)<br>(87%) | HIS878[60, 61]<br>ASP1031 | HIS1213<br>ASP1208 | 0.32 | 4.31 |
| TE/MPT<br>(2215-2500/1311-1709)<br>(72%) | SER2308[38]<br>ASP2338<br>HIS2841 | SER1417<br>HIS1545 | - | - |
| ER/ER<br>(1527-1863/367-871)<br>(67%) | LYS1771[11]<br>ASP1797 | HIS553<br>THR616 | - | - |

\* The $Q_H$ value is a metric for structural homology \*\* adjusted based on our simulation trajectory

### 2.4 Identifying coordinating and pocket waters

Coordinating waters that formed hydrogen bonds with catalytic residues were identified using an inhouse TCL script, and visualized using H-bond tool in VMD. A distance cutoff of 3 Åand an angle cutoff of 20^°^ were used. Pocket waters were identified using a cutoff distance of 3 Åfrom the pocket grid-points obtained from traj_cavity.

## 3 RESULTS AND DISCUSSION

### 3.1 Stability of simulated FAS-I complexes and their domains

The stability of hFAS-I and MtbFAS-I complexes were evaluated using root-mean-square deviation (RMSD) of backbone atoms of individual chains. The mean and standard deviation of the RMSD was computed using the last 25 ns of the 100 ns-long simulation trajectories. In the hFAS-I complex, the backbone RMSD of the two chains A and B in hFAS-I, without excluding any residues, was found to be large, with mean RMSD of 16.8 Å and 20.5 Å (data not shown). hFAS-I has several flexible linkers and a mobile TE domain owing to its relatively open structure. It also has two additional pseudo-domains (*ψ*ME and *ψ*KR) between DH and ER. Figure 1a shows the backbone RMSD of the two chains in hFAS-I upon excluding all residues corresponding to TE, linkers and the pseudo-domains. The mean RMSD of the two chains based on the remaining domains was observed to be 7.6 Å and 6.6 Å, respectively. Upon additionally excluding residues corresponding to ACP, the RMSD of Chain A (7.6 Å) remained nearly the same as above, while the RMSD of chain B (5.8 Å) further decreased (data not shown). The backbone RMSD of the six MtbFAS-I chains (A-F), without excluding any domains, is shown in Figure 1b. Individual chains in both complexes fluctuated around stable RMSD values in the last 25 ns of respective simulation trajectories, indicating stability of the overall complexes in both trajectories.

**FIGURE 1.**
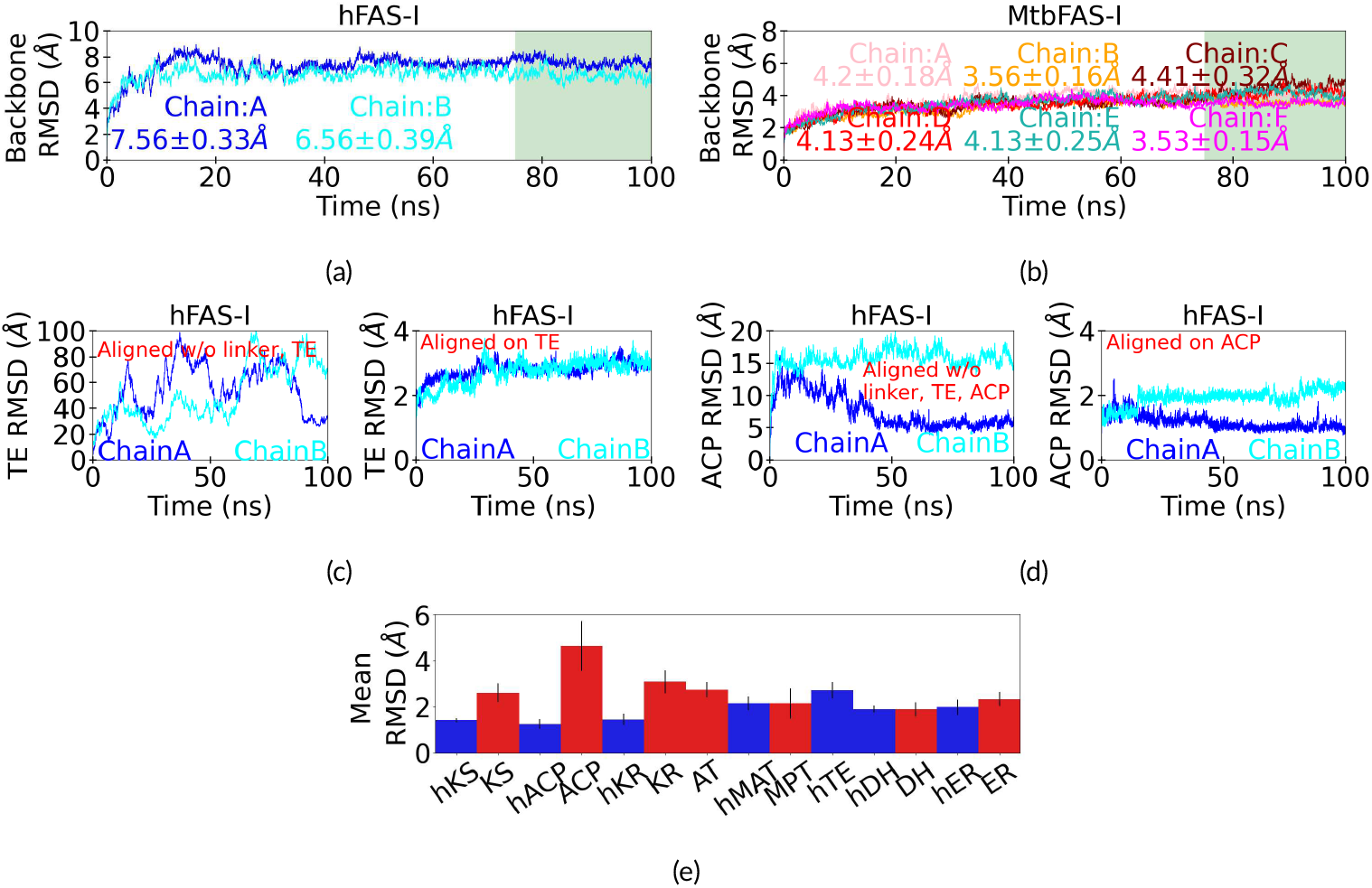
Stability of the hFAS-I and MtbFAS-I and their domains. Time evolution of (a) backbone RMSD of hFAS-I aligned to backbone excluding TE, pesudodomain and linkers and (b) backbone RMSD of MtbFAS-I aligned to its backbone. Legends display mean ± standard deviation in the last 25 ns (green region) for all chains. (c-d) TE and ACP Backbone RMSD when backbone aligned to backbone excluding TE and linkers and when TE and ACP aligned to itself. shows the high mobility of TE in hFAS-I.(e) Shows domain-wise comparison of mean RMSD between hFAS-I and mtbFAS-I. Error bars show the standard deviation in this and the subsequent figures

The exclusion of TE and linker residues in calculating the RMSD of hFAS-I chains resulted in lower RMSD. To investigate this further, we visualized the trajectory in VMD [62] to discover that significant motion of TE domain relative to the rest of the complex throughout the 100 ns trajectory was the largest contributor to fluctuations in the structure of the complex. We calculated the RMSD of TE using two different methods of aligning the trajectory. The left panel in Figure 1c shows the RMSD of TE calculated after aligning the trajectory using backbone atoms of the chain excluding the TE, linkers and the pseudo-domains. RMSD of TE, calculated after aligning backbone atoms of TE, is shown in the right panel of Figure 1c. A comparison of the RMSD using the two alignment methods reveals that although TE domain fluctuates to a significant extent relative to rest of the chain, as seen from the relatively high and significantly fluctuating RMSD in the left panel, its internal structure is relatively rigid, with RMSD remaining well below 4 Å throughout the simulation. In agreement with previous reports, we observe that TE is a mobile domain that undergoes significant rigid-body translation and rotation at the simulated timescale of 100 ns[30]. The RMSD of ACP was calculated using the two alignment methods, first calculating ACP’s RMSD after the entire chain, excluding linker, TE and ACP, was aligned; second, ACP’s RMSD was calculated after ACP aligned onto itself shown in Figure 1d. Although not to the same extent as TE, ACP moderately fluctuates relative to the rest of the chain (left panel, alignment excludes ACP, TE, linkers and pseudo-domains), while maintaining its internal rigid structure (right panel, alignment on ACP).

As summarized in the discussion on the mechanism of fatty acid synthesis in the section Introduction, the fatty acid elongation stage involves shuttling of the growing substrate by ACP between KS, KR, DH and ER. These domains are common in hFAS-I and MtbFAS-I. In hFAS-I, MAT transfers acetyl and malonyl moieties to KS in the initiation stage, and malonyl moiety alone during the elongation stage, while a distinct TE domain terminates the synthesis. In MtbFAS-I, AT transfers acetyl moiety in the initiation stage, while MPT transfers malonyl moiety during the initiation and elongation stages and terminates the synthesis. Thus, hFAS-I MAT’s function partly resembles the functions of both AT and MPT in MtbFAS-I. MtbFAS-I MPT’s function additionally resembles the function of TE in hFAS-I. Column 1 in Table 1 lists corresponding domains between the two complexes based on their roles in the mechanism of fatty acid synthesis. Number ranges in brackets indicate the range of residues assigned to each domain in the two species and the percentage of residues in hFAS-I domain relative to the corresponding MtbFAS-I domain. ACP, KS, KR, DH and ER are significantly shorter (based on residues count) in hFAS-I. TE is shorter than the corresponding MPT. In contrast, hFAS-I MAT is marginally longer than AT and almost as long as MPT in MtbFAS-I.

The RMSD of each domain in each of the two complexes was calculated by aligning the backbone atoms of the domain of each chain to the first frame in the trajectory. Figure 1e shows the mean RMSD of each domain in hFAS-I and MtbFAS-I, averaged over the last 25 ns of the trajectory and averaged over all the chains in the complex (i.e. averaged over two chains in hFAS-I, averaged over six chains in MtbFAS-I). Corresponding domains between the two species (see Table 1) are shown adjacent to each other in Figure 1e. With the exception of MtbFAS-I ACP, the mean RMSD of all domains is below 3 Å. The mean RMSD of KS, ACP, KR, and ER is higher in MtbFAS-I than in hFAS-I. DH in both species shows a comparable mean RMSD, as do the corresponding domains MAT and MPT. In comparison to MAT, AT in MtbFAS-I is observed to be more flexible, while TE is more flexible than the (partly) functionally resembling MPT. hFAS-I has a short ACP that is missing the structural core present in its MtbFAS-I counterpart. Thus, nearly all domains are less or equally flexible in hFAS-I, with the exception of TE. Further, figure 1e shows that all seven domains in hFAS-I and in MtbFAS-I are observed to exhibit a stable internal structure.

### 3.2 Structural comparison of domains and fluctuations of mapped residues

Despite differences in the sequence length and dynamics of corresponding domains between hFAS-I and MtbFAS-I, we observed a remarkable structural similarity among several domains using STAMP structural alignment. The following pairs of domains, one from hFAS-I and the corresponding second domain from MtbFAS-I, were successfully aligned: MAT/AT, MAT/MPT, KS/KS, ACP/ACP, KR/KR, and DH/DH (see Table 1). The TE/MPT and ER/ER domain pairs could not be aligned using STAMP structural alignment. Figure 2 shows the aligned domains, with black color representing the highest level of similarity and blue (hFAS-I) or red (MtbFAS-I) colors representing the lowest levels of similarity.

**FIGURE 2.**
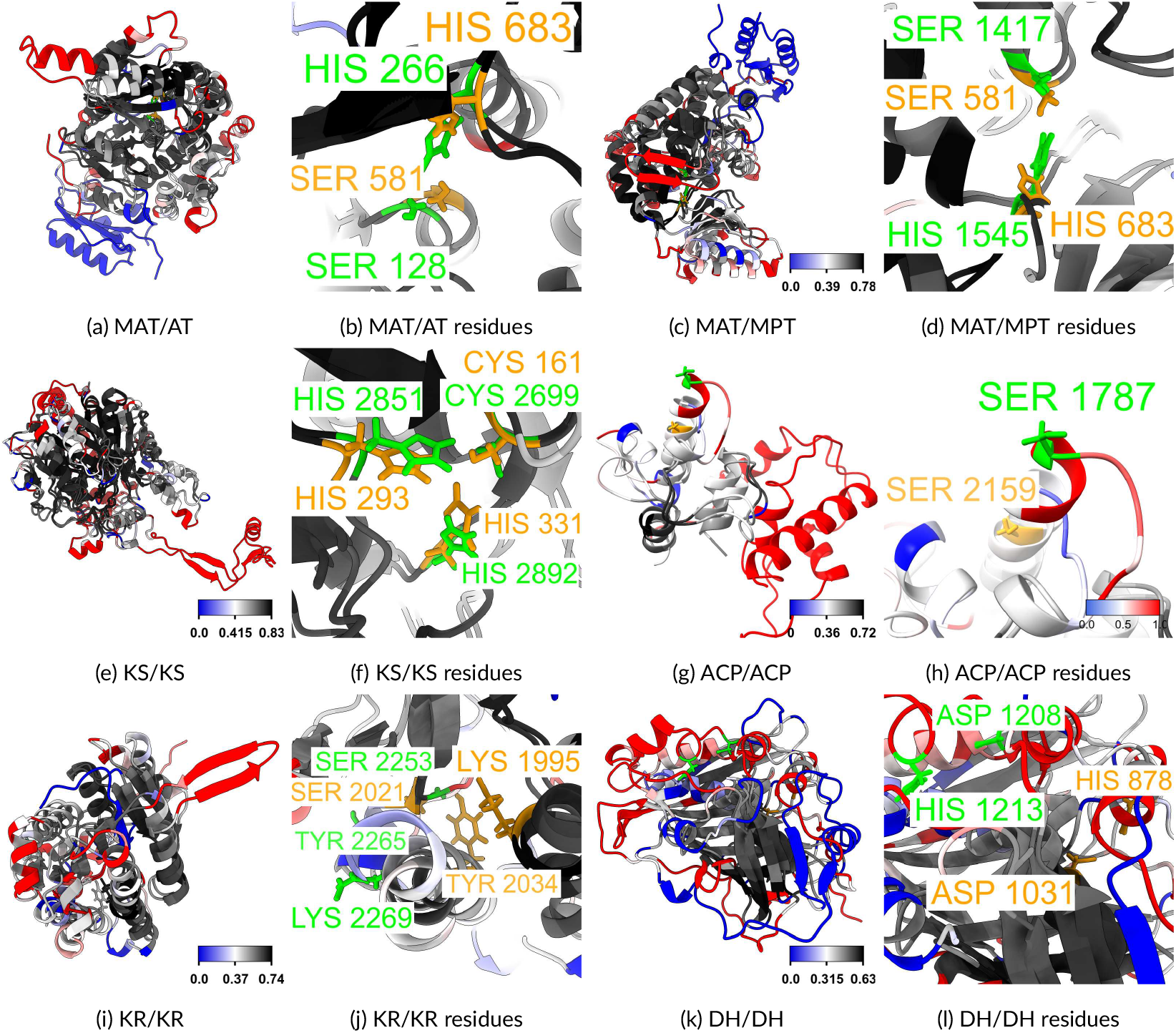
Panels (a)-(l) depict the STAMP structural alignment of each domain in hFAS-I and MtbFAS-I, along with a magnified view showcasing their catalytic residues represented in stick form. Darker shades of grey represent regions with highest structural similarity, while darker shades of blue color (hFAS-I) or red color (MtbFAS-I) represent regions with increasingly dissimilar structures.

#### 3.2.1 hFAS-I MAT, MtbFAS-I AT

The highest structural homology score Q_*H*_ is found between hFAS-I MAT and MtbFAS-I AT (Table 1). STAMP alignment of these domains, which also play a partially similar role in the mechanism of fatty acid synthesis, is shown in Figure 2a. Structural similarity is represented at the residue level using a color scale based on Q_*res*_ scores of residues, with black color representing the highest Q_*res*_ score and hence the most similar residues, and either blue colour (for hFAS-I) or red colour (for MtbFAS-I) representing a zero Q_*res*_ score and hence the most dissimilar residues. The colour scale for hFAS-I is shown at the bottom of the figure. hFAS-I MAT is a longer domain in comparison to MtbFAS-I AT. Excess residues in hFAS-I MAT are primarily seen in alpha helices and a beta-sheet in blue colour at the bottom of the figure.

MtbFAS-I AT also has a unique alpha helix that is absent in hFAS-I MAT, appearing in red colour at the top of the figure. Apart from these major regions and a few minor regions, the rest of the structure have a significant extent of similarity between the two species, seen as regions in bold shades of grey.

To compare the dynamics of structurally similar residues, we found all the residues with a Q_*res*_ score of 0.5 or above. The resulting 225 residues are serially listed in Table S1. Figure 3a shows the RMSF of these residues in the two complexes. The RMSF of these structurally similar residues with a high Q_*res*_ seem to be comparable in the two domains, with the exception of a few residues. Large differences in RMSF are seen for regions corresponding to a region with unstructured secondary structure (except THR333) in vicinity of residues labelled in cyan and magenta colors, *viz*. THR541 and GLY679 in hFAS-I, and HIS1 and THR333 in MtbFAS-I. Overall, corresponding resiudes in both these domains do not show any large fluctuations.

**FIGURE 3.**
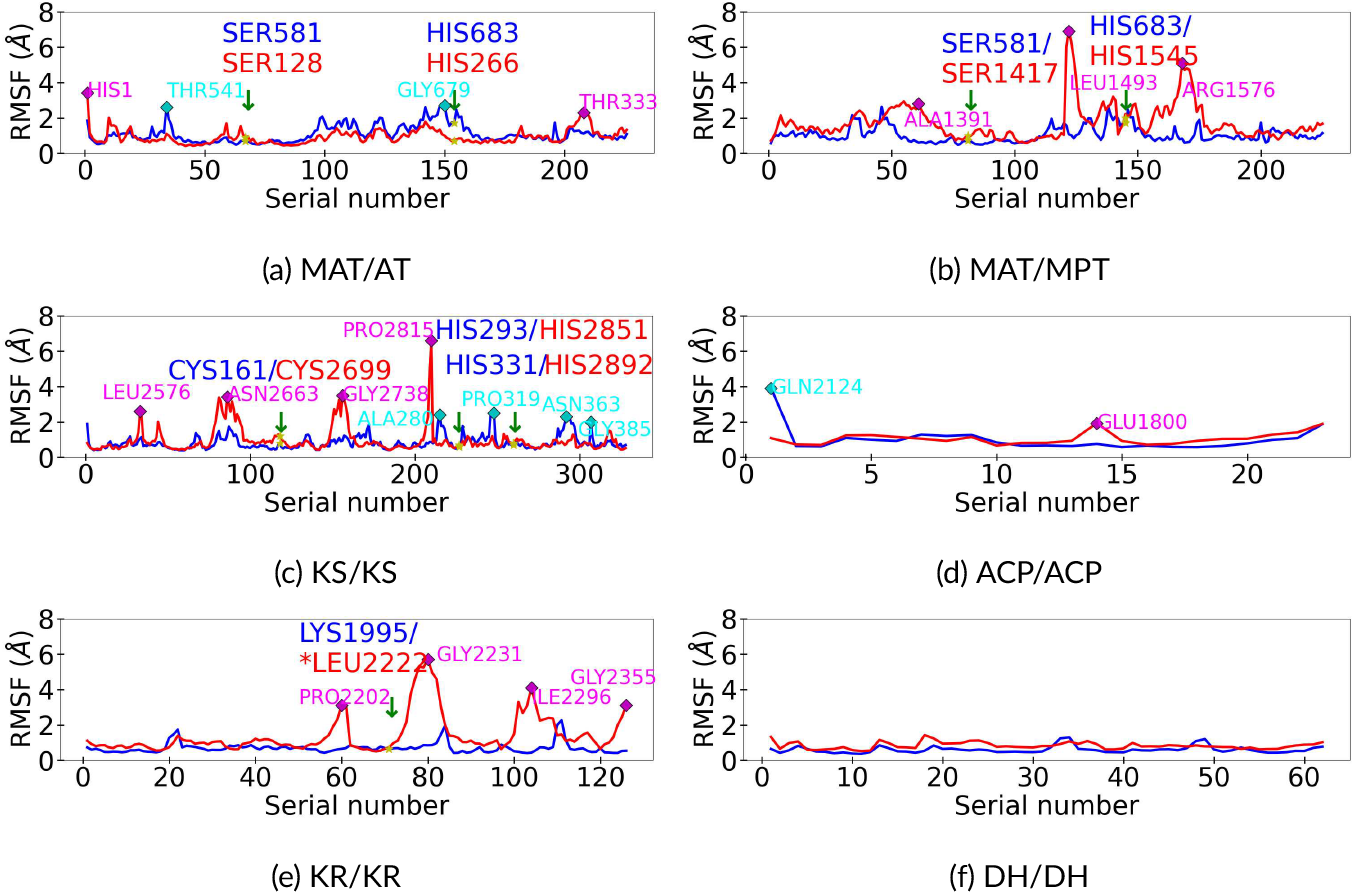
Domain-level flexibility of mapped residues with Q_*res*_ > 0.5 in hFAS-I (blue) and MtbFAS-I (red). Residue corresponding serial numbers are listed in Tables S1-S6 of Supporting Information. Catalytic residues in hFAS-I/MtbFAS-I are labelled in blue/red color and marked with arrows. hFAS-I/MtbFAS-I residues corresponding to peak RMSF values are labelled in cyan/magenta color and marked as diamonds.

Figure 2b shows a magnified view of the aligned structures with a detailed view of the catalytic residues, *viz*. SER581 and HIS683 in hFAS-I [52], and the corresponding SER128 and HIS266 in MtbFAS-I [6]. Sidechains of these residues are shown using a stick representation, with orange color representing sidechains of hFAS-I and green color representing sidechains of MtbFAS-I. Although hFAS-I MAT and MtbFAS-I AT have different lengths and dynamics, orientation of these catalytic residues is found to be similar. The RMSF of these resiudes is also found to be low and comparable, as seen from Figure 3a. Thus, although both MAT and AT show unique secondary structures near their surfaces, they have a remarkable structural similarity in the region around the catalytic residues.

#### 3.2.2 hFAS-I MAT, MtbFAS-I MPT

In addition to MtbFAS-I AT, hFAS-I MAT also shares a common mechanistic role with MtbFAS-I MPT. MAT and MPT also show the second highest structural homology score Q_*H*_ = 0.47 (Table 1) and show a comparable mean RMSD (Figure 1e). These domains are also of a comparable length. STAMP alignment of these domains is shown in Figure 2c. Black regions denote high structural similarity signified by large Q_*res*_, while blue color (hFAS-I) and red color (MtbFAS-I) denote low structural similarity. Color scale for hFAS-I is shown at the bottom. We have observed two distinct regions: the region in hFAS-I with multiple helices seen in blue color at the top-right of the figure, and the region in MtbFAS-I with a beta sheet and helices seen in red color in the mid-region of the figure. Rest of the residues in the domain show high structural similarity.

The 222 residues with Q_*res*_ ≥ 0.5 are serially listed in Table S1. Figure 3a shows the RMSF of these residues in the two complexes. The RMSF of these structurally similar residues seem to be similar, except for regions with unstructured secondary structure in vicinity of residues labelled in cyan and magenta colors. Specifically, the two regions around LEU1493 and ARG1576 in MtbFAS-I exhibit large relative fluctuations. These regions, however, lack a rigid secondary structure and appear as loops. The catalytic residues SER and HIS show comparable RMSF in the two complexes. Thus, the catalytic regions of the two domains have a highly similar structure and dynamics.

Figure 2d provides a magnified view of the aligned structures, showing the orientation of side-chains of the catalytic residues. The catalytic residues from MAT (SER-581 and HIS-683) and MPT (SER-1417 and HIS-1545) exhibit remarkably similar orientations in the aligned domains.

#### 3.2.3 hFAS-I and MtbFAS-I KS

The functionally identical KS domain in the two FAS-I complexes also shows a large structural similarity, indicated by its high Q_*H*_ score in Table 1. The STAMP alignment of the two KS domains is depicted in Figure 2e, using a coloring scheme similar to that in the previous sections. KS is longer in MtbFAS-I compared to hFAS-I, with excess residues primarily belonging to the extended region at the bottom of the figure shown in red color. This extended region, consisting of a beta sheet and a helix, is a part of the wheel in MtbFAS-I complex. A few other structurally distinct residues appear as red helices in the top-left of the figure.

Based on a threshold value of 0.5 for the Q_*res*_ score, we found 328 mapped residues with high structural similarity. These residues are serially listed in Table S3. The RMSF of these residues in both complexes is depicted in Figure 3c. With the exception of the regions around residues labelled in cyan and magenta colors, *viz*. ALA280, PRO319, ASN363, GLY385 in hFAS-I and LEU2576, ASN2663, GLY2738, PRO2815 in MtbFAS-I KS, the RMSF of other structurally similar residues appears to be low and comparable in both complexes. The residues listed above, corresponding to regions with large RMSF, do not adopt a rigid secondary structure, with the exception of the short helix forming residue GLY2730.

Figure 2f provides a magnified view of the aligned structures, showing the side-chain orientation of the catalytic residues: CYS161, HIS293 and HIS331 in hFAS-I, and CYS-2699, HIS-2851, and HIS-2892 in MtbFAS-I. Sidechains are represented in a stick representation, with orange indicating hFAS-I and green representing MtbFAS-I. Despite the differing lengths of hFAS-I MAT and MtbFAS-I AT, the orientation of these catalytic residues is found to be identical, suggesting that despite the large differences in the length and structure of the two KS domains, the region in the vicinity of their catalytic core is nearly identical. Further, fluctuations of catalyic residues are also comparable between the two KS domains (Figure 3c).

#### 3.2.4 hFAS-I and MtbFAS-I ACP

Although the fluctuation function of ACP in both FAS-I systems is similar, hFAS-I ACP is a relatively short domain consisting of a catalytic core, while MtbFAS-I ACP includes an additional structural core composed of four *α* -helices. Figure 2g shows the STAMP alignment of ACP in both complexes. Almost the entire hFAS-I is mapped onto the half of MtbFAS-I ACP that constitutes its catalytic core. Of the four *α* -helices in the catalytic core, only one shows a high degree of structural similarity. An additional loop region that connects helix 1 to helix 2, highlighted in black, exhibits a high degree of structural similarity. Most of the remaining residues, including those forming the three additional *α* - helices shaded in gray color with a low Q_*res*_ score have low degree of structural similarity. MtbFAS-I ACP’s recognition helix also contains a few extra amino acids. These appear in red color at the top-left of the figure.

Using a threshold of Q_*res*_ ≥ 0.5, we have found 23 mapped residues that are listed in Table S4. RMSF of these residues are shown in Figure 3d. Fluctuations of these mapped residues are comparable. A moderate deviation is observed between the RMSF values of hFAS-I residue THR2165, which present in a loop structure, and the corresponding MtbFAS-I residue GLU1800 located at the end of the recognition helix Figure 3d.

Figure 2h offers a magnified view of the catalytic core of the two ACPs, featuring the conserved SER residue (SER-1787 in MtbFAS-I and SER2195 in hFAS-I) using a stick representation. This conserved SER residue acts as the site where the growing fatty acid substrate attaches [23, 63, 64, 65]. Interestingly, these conserved residues are not located at an identical position in the two enzyme complexes. Despite the recognition helix having a similar length of 15 residues in both ACPs, SER2195 is positioned four residues away from the end of the helix in hFAS-I, whereas in MtbFAS-I, the corresponding SER1787 residue is situated at the end of the helix. Additionally, *α* -3 helix in hFAS-I consists of 6 residues (VAL2176 to GLN2181), whereas the same helix consists of 4 residues (SER1812 to LEU1814) in the MtbFAS-I ACP average structure from our simulation.

#### 3.2.5 hFAS-I and MtbFAS-I KR

Similar to ACP, KR is shorter in hFAS-I in comparison with MtbFAS-I (Table 1). Despite the length difference, KR from the two complexes shows a significant structural similarity as indicated by the high Q_*H*_ score. Figure 2i displays the STAMP alignment of KR from both FAS-I systems. MtbFAS-I is seen to be larger in size compared to its hFAS-I counterpart, with an additional extended *β* -sheet present in MtbFAS-I, seen in red color at the top-right of the figure. Additionally, an extra alpha helix and a loop region shown in red color are seen in MtbFAS-I. Despite these differences, several residues in the interior of the domain show higher structural similarity shown with increasingly dark shades of grey.

Using the same threshold value of Q_*res*_ ≥ 0.5, we found 126 mapped residues with high structural similarity in KR. These residues are listed in Table S5. RMSF of these residues are shown in Figure 3e. We observed that in the regions around residues PRO2202, GLY2231, LEU2296 and GLY2355 in MtbFAS-I KR displayed higher fluctuations compared to hFAS-I KR. All these residues, with the exception of GLY2231, are part of unstructured loop regions. GLY2231 is present in the middle of a long helix, and yet exhibits large fluctuations.

Figure 2j shows a magnified view of the catalytic residues of hFAS-I (LYS1995, SER2021, TYR2034) and MtbFAS-I (SER2253, TYR2265, LYS2269). The corresponding pairs of catalytic residues of KR in the two complexes are in the same region, but unlike in many other domains, they do not align well.

#### 3.2.6 DH: distinct domain

STAMP analysis of the DH domains in hFAS-I and in MtbFAS-I revealed lowest structural similarity among the other domains, as seen from the low Q_*H*_ score of 0.32 (Table 1). Figure 2k depicts the alignment of DH from the two enzymatic complexes using a color scale based on Q_*res*_ score. Darker shades of grey primarily concentrated in the interior of the domain denote increasingly higher structural similarity in this region. This structurally similar region largely consists of residues forming beta sheets. Regions closer to the domain surface are increasingly dissimilar between the two complexes. Several structures in blue color denote structurally distinct regions in hFAS-I, while those in red color denote regions with structurally most dissimilar residues in MtbFAS-I. These include residues belonging to unstructured loops, helices and beta sheets that are located near the peripheral region of DH.

We identified only 62 DH residues with high structural similarity (Q_*res*_ ≥ 0.5) in the two complexes. These residues are listed in Table S6. The RMSF analysis of these 62 residues reveals comparable fluctuations, all of which are well below 2 Å, as illustrated in Figure 3f. These low fluctuations can be attributed to the fact the many of these residues form *β* -sheets.

Among all the corresponding domains discussed above, hFAS-I DH exhibits notable structural differences relative to MtbFAS-I DH, with only a small fraction of residues showing a high structural similarity. Thus, we identified DH as one of the most distinct domains and investigated its structure for catalytic pocket identification and analysis.

Figure 4a shows a combined view of the STAMP aligned hFAS-I and MtbFAS-I DH domains and their catalytic pockets. The catalytic pockets are shown using surface representation, with pink and yellow surface views showing the catalytic pockets in DH from hFAS-I and MtbFAS-I, respectively. Figure 4b provides a detailed view of the catalytic pocket in hFAS-I DH. This pocket includes the catalytic residues HIS787 and ASP1031, shown using a stick representation. Similarly, the catalytic pocket and catalytic residues participating in its formation in MtbFAS-I DH are shown in Figure 4c.

**FIGURE 4.**
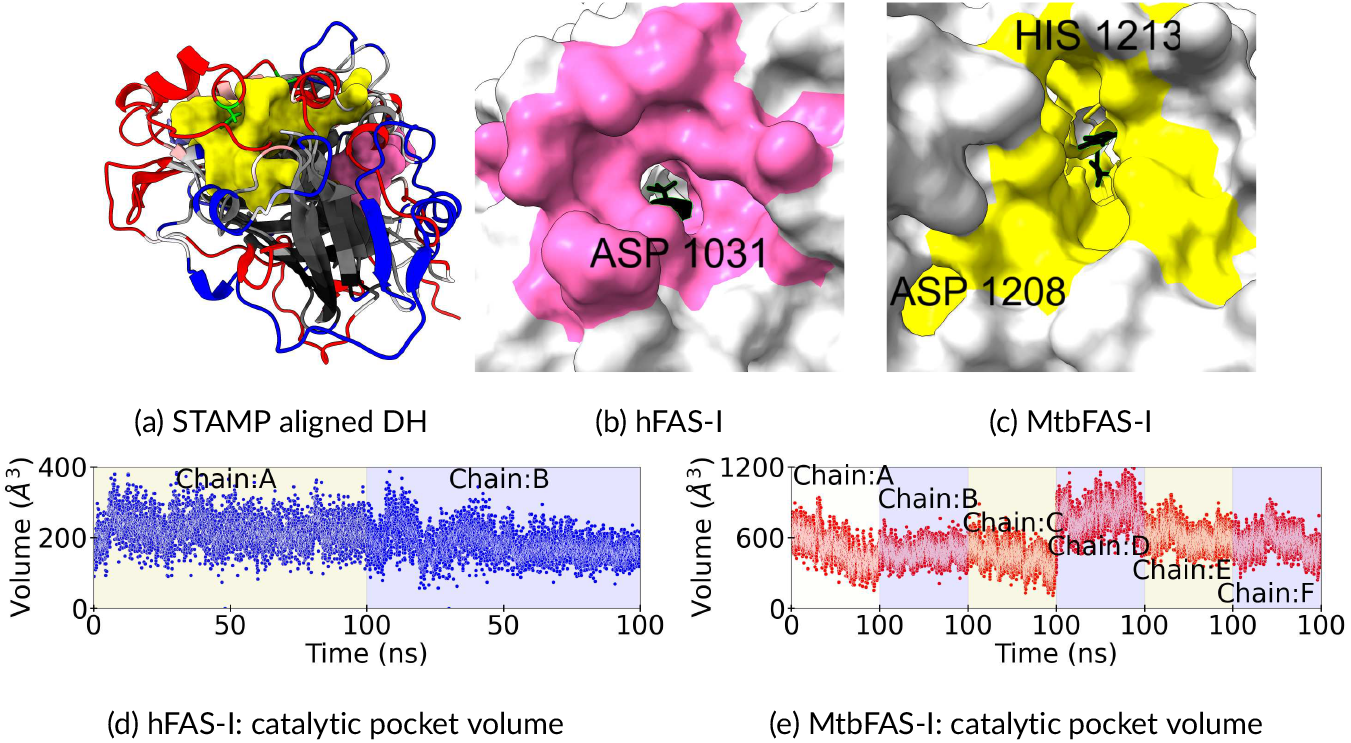
(a) Aligned hFAS-I and MtbFAS-I DH domains with their catalytic pocket shown in pink and yellow surface respectively. (b) DH of hFAS-I. (c) DH of MtbFAS-I (d) Variation of hFAS-I DH’s catalytic pocket volume in the 100 ns trajectory. (e) Variation of MtbFAS-I DH’s catalytic pocket volume in the 100 ns trajectory.

As described in Methods, of the multiple pockets predicted for the average domain structure, the largest pocket containing catalytic residues was chosen to be the catalytic pocket. The catalytic pockets identified for DH in hFAS-I and MtbFAS-I were predicted as the largest and second largest pockets, respectively. In hFAS-I, the catalytic pocket is formed by 32 residues, while 68 residues form the pocket in MtbFAS-I. Analyzing our hFAS-I simulation trajectory for dynamics of these pockets gives a mean volume of 200.9±48.0 Å^3^, with a variation in the range from 57.6 Å^3^ to 408.9 Å^3^ in the 100 ns trajectory. We, have also observed that volume was zero in two instances, i.e., the pocket was closed in two frames. In the MtbFAS-I trajectory, the mean volume is 567.4±153.5 Å^3^, which varies from 112.5 Å^3^ to 1193.64 Å^3^. These data show that the pocket in MtbFAS-I DH is substantially larger and more flexible compared to hFAS-I DH.

A comparison of the physicochemical properties of these pockets is summarized in Table 2. DH’s catalytic pocket in hFAS-I is less hydrophobic and more polar compared to MtbFAS-I. However, hFAS-I contains comparable aromatic residues. The percentage of positive and negative charges in Mtb DH are similar, whereas, in hFAS-I, the percentage of the negative charges is higher than the number of positive charges.

**TABLE 2.** Residues forming a catalytic pocket in the DH and ER domains of hFAS-I and MtbFAS-I and their characteristics.

| Properties | hFAS-I DH | MtbFAS-I DH | hFAS-I ER | MtbFAS-I ER |
| --- | --- | --- | --- | --- |
| Total count | 32 | 68 | 75 | 110 |
| Hydrophobic residues | 53.1% | 57.4% | 52.0% | 52.7% |
| Polar residues | 18.1% | 8.8% | 18.6% | 26.4% |
| Negative residues | 9.4% | 10.3% | 5.3% | 10.0% |
| Positive residues | 6.3% | 11.7% | 13.3% | 10% |
| Aromatic residues | 12.5% | 11.8% | 10.7% | 0.9% |

Catalytic residues ASP74 and HIS70 have been identified in Escherichia coli *β* -hydroxydecanoyl thiol ester dehydrate that is involved in dehydration and isomerization of the growing substrate. [66]. In their crystal structure, these authors reported a water molecule coordinating with these two catalytic residues. Corresponding catalytic residues ASP1033 and HIS1037 in mammalian FAS-I are involved in the dehydration reaction mechanism that results into release of a water molecule. [13, 67]. A possible dehydration reaction mechanism would involve coordinating of this water molecule with both the catalytic residues before being released into the bulk. In our Mtb trajectory, we find several frames where hydrogen from a water molecule forms a persistent hydrogen bond with oxygen of ASP1208. On several instances, we also observe the same water forming an additional hydrogen bond with HIS1213 in one of the following ways: either between its oxygen and hydrogen on NE2 of HIS1213, or between its hydrogen and the nitrogen ND1 of HIS1213 (Supplementary Figure S3).

We have also computed the number of water molecules in the volume spanned by the catalytic pocket in each frame and in each chain. In our MtbFAS-I trajectory, DH’s catalytic pocket was found to have 53 ± 7 water molecules on an average. In contrast, the average number of water molecules in the catalytic pocket of hFAS-I DH was merely 16 ± 3. Noting that the average volumes of the two pockets are significantly different, there are more water molecules per volume of the pocket in MtbFAS-I DH compared to hFAS-I.

#### 3.2.7 ER: distinct domain

We were not able to successfully employ the STAMP algorithm to structurally align ER from the two FAS-I complexes, even with the lowest constraints implemented in VMD[62] (scanscore = 0, scanslide=1). After several iterations, we were able to obtain a partially similar structure, as follows. We performed a sequence similarity using ClustalW algorithm implemented in VMD [62], and employed STAMP algorithm on the longest section of 75 sequentially similar residues (hFAS-I residues CYS1759 to GLY1821, MtbFAS-I residues SER670 to SER732). This resulted into a partial structural alignment of ER based on the selected residues (Figure 5a). We note that this choice of a subset of ER residues for partial STAMP structural alignment is not unique. Using other sequentially similar residues did not result into a significantly better overlap of the two ER domains than the partial overlap seen in the figure.

**FIGURE 5.**
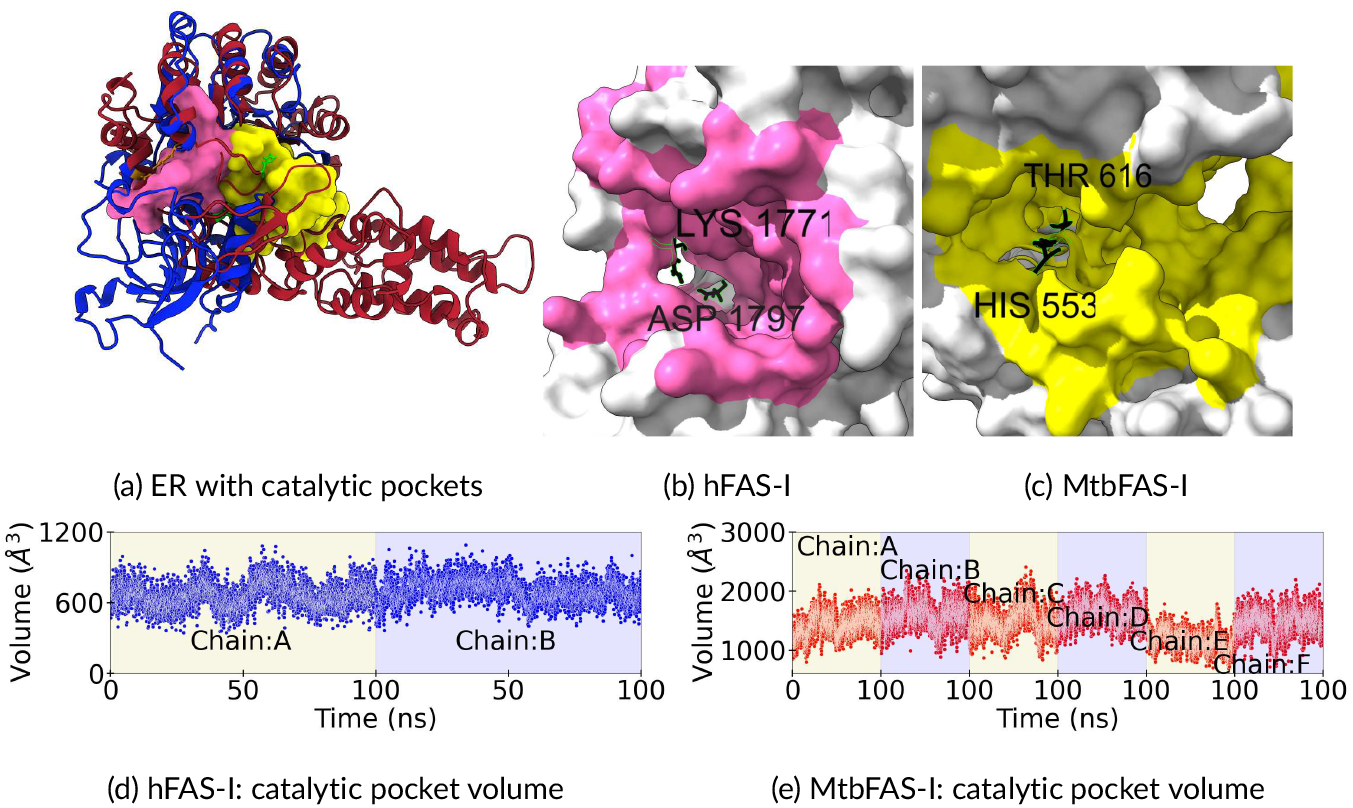
(a) Aligned hFAS-I and MtbFAS-I ER domains with their catalytic pocket shown in pink and yellow surface respectively. (b) ER of hFAS-I. (c) ER of MtbFAS-I (d) Variation of hFAS-I ER’s catalytic pocket volume in the 100 ns trajectory. (e) Variation of MtbFAS-I ER’s catalytic pocket volume in the 100 ns trajectory.

Due to lack of structural alignment in a larger fraction of the domain, we could not identify any structurally similar residues between the two ERs using a Q_*res*_ score. Further, the catalytic residues in hFAS-I ER (LYS-1771, ASP-1797) are of a different nature than in MtbFAS-I ER (HIS-553, THR616) despite their similar roles in the mechanism of fatty acid synthesis, which is a unique feature of ER that is not observed in other domains.

A combined view of the catalytic pockets of ER in the two complexes is also shown in Figure 5a. Figure 5b displays the catalytic pocket in ER of hFAS-I using a surface representation. Figure 5c showcases the ER domain of MtbFAS-I, with blue surface residues indicating the catalytic pocket. In hFAS-I ER, the catalytic pocket is formed by 75 residues, while 110 residues form the pocket in MtbFAS-I ER. In our 100 ns simulation trajectory, the hFAS-I ER pocket exhibits a mean volume of 681.1±108.8 Å^3^ and ranges from 345.7 Å^3^ to 1089.4 Å^3^. The catalytic pocket of ER in MtbFAS-I shows a mean volume of 1460.8±240.26.9 Å^3^, with a range from 694.2 Å^3^ to 2406.5 Å^3^. It is interesting to note that a significantly large pocket is present in MtbFAS-I ER in comparison to hFAS-I ER. We have verified that the catalytic pocket that we have identified in MtbFAS-I ER is not blocked by either its binding cofactor flavin mononucleotide or by other domains in the complex (Figures S1, S2).

The ER catalytic pocket in hFAS-I and MtbFAS-I showed similar hydrophobic percentages. Unlike in DH, the pocket of hFAS-I ER has a much higher percentage of aromatic residues compared to the pocket of MtbFAS-I ER. The pocket of MtbFAS-I ER has more negatively charged residues compared to the pocket of hFAS-I ER. hFAS-I ER’s pocket has slightly larger percentage of positively charged residues than MtbFAS-I. However, MtbFAS-I ER’s pocket has more number of positive residues owing to its larger size. We observed that the percentage of positive and negative charges in MtbFAS-I ER’s pocket, like DH, are balanced, whereas in hFAS-I, the percentage of positive charges is double the percentage of negative charges.

The catalytic pocket in MtbFAS-I ER was found to have an average of 132 ± 13 water molecules in our trajectory. In contrast, our hFAS-I ER was found to have an average of 74 ± 8 water molecules. Despite the large pocket volume in MtbFAS-I ER, it has a significantly smaller number of water molecules per volume compared to hFAS-I ER.

## 4 CONCLUSION

In conclusion, we have found that ER is strcuturally the most distinct domain in MtbFAS-I with reference to hFAS-I, followed by DH. The catalytic pockets identified in DH and ER are larger in volume and more flexible in MtbFAS-I. Thus, steric interations should play a major role in selectivity of a ligand towards MtbFAS-I’s ER. hFAS-I has a higher number of aromatic residues compared to MtbFAS-I. Therefore, *π*-*π* interactions should be discounted while designing ligands. The ER domain of MtbFAS-I is 50% longer and its catalytic pocket is 45% larger in comparison to hFAS-I ER. Thus, although the percentage of aliphatic hydrophobic residues is similar in both the pockets, the larger pocket in MtbFAS-I implies that it has more hydrophobic residues. The pocket is net positively charged in hFAS-I, but is neutral in MtbFAS-I. The pocket in MtbFAS-I has more polar residues. These differences can also be exploited to design ligands selectively targeting MtbFAS-I’s ER.

## Acknowledgements

This research was supported in part by Science and Engineering Research Board (SERB) through the Start-up Research Grant Number SRG/2020/002186 and in part by Indian Institute of Technology Kanpur through the Initiation Grant. The computational support and the resources provided by PARAM Sanganak under the National Supercomputing Mission, Government of India at the Indian Institute of Technology Kanpur are gratefully acknowledged. A.K. also acknowledges financial support from Institute Postdoctoral Fellowship at Indian Institute of Technology Kanpur.

## Data Availability

The data that support the findings of this study are available from the corresponding author upon reasonable request.

## Supporting Information

Supporting Information (Document S1) can be found online.

- Document S1: Figure S1: flavin mononucleotide and ER’s catalytic pocket in MTBFAS-I; Figure S2: catalytic pockets in DH and ER with ACP in MTBFAS-I; Figure S3: Simulation snapshots showing water coordinating with catalytic residues in DH Table 1: hFAS-I MAT and MtbFAS-I AT mapped residues table;Table 2: hFAS-I MAT and MtbFAS-I MPT mapped residues table; Table 3: hFAS-I KS and MtbFAS-I KS mapped residues table; Table 4: hFAS-I ACP and MtbFAS-I ACP mapped residues table; Table 5: hFAS-I KR and MtbFAS-I KR mapped residues table;Table 6: hFAS-I DH and MtbFAS-I DH mapped residues table

