## Supplementary Document S1 for "Unraveling Structural Disparities in Human and Mycobacterium Tuberculosis Type-I Fatty Acid Synthase"

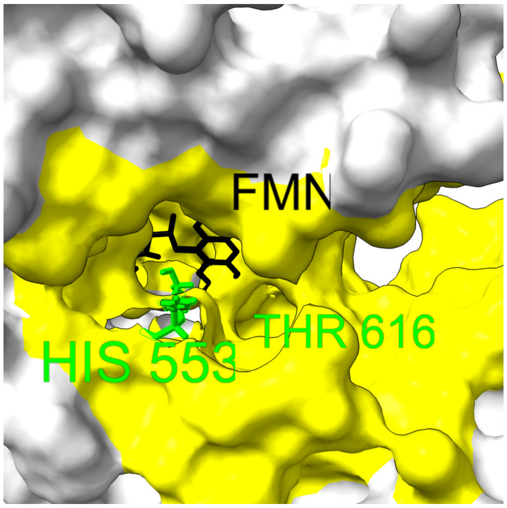


Figure S1: The figure depicts the entrance of the MtbFAS-I ER's pocket along with flavin mononucleotide (FMN), demonstrating that the pocket remains open and FMN does not obstruct it. The alignment of chain A from PDB ID: 6JGC on the average structure from our simulation indicates the FMN location. The grey and yellow surfaces represent the ER domain and the ER pocket-forming residues, respectively. The lime and black sticks correspond to the MtbFAS-I ER's catalytic residues and FMN, respectively.

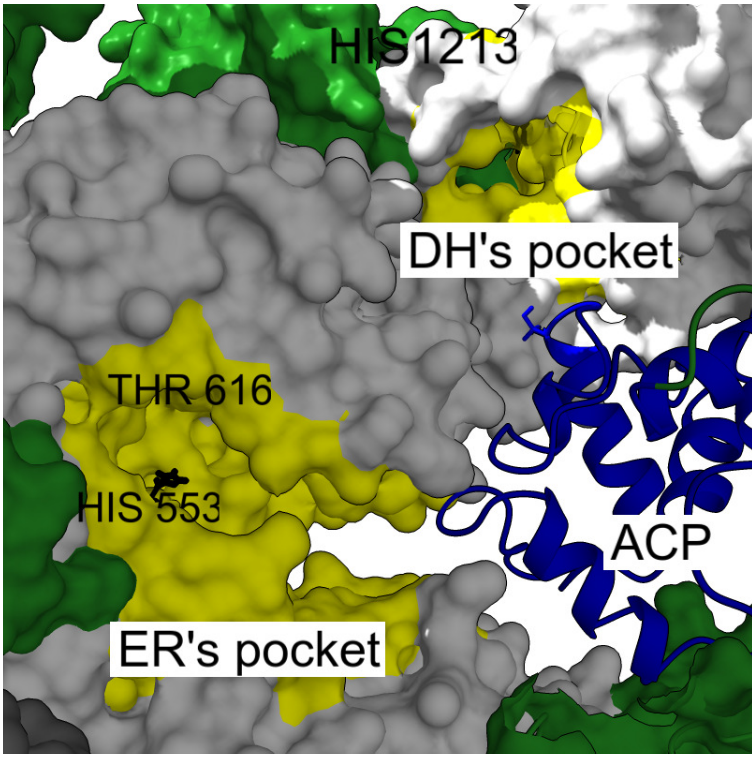


Figure S2: Figure illustrating the ER and DH domain pocket highlighted in yellow surface, showing it is unobstructed by any other domains from its own chain (green surface) or neighbouring chains(grey surface). ACP (orange cartoon) stalled at DH, indicating the entrance of the pocket at DH.

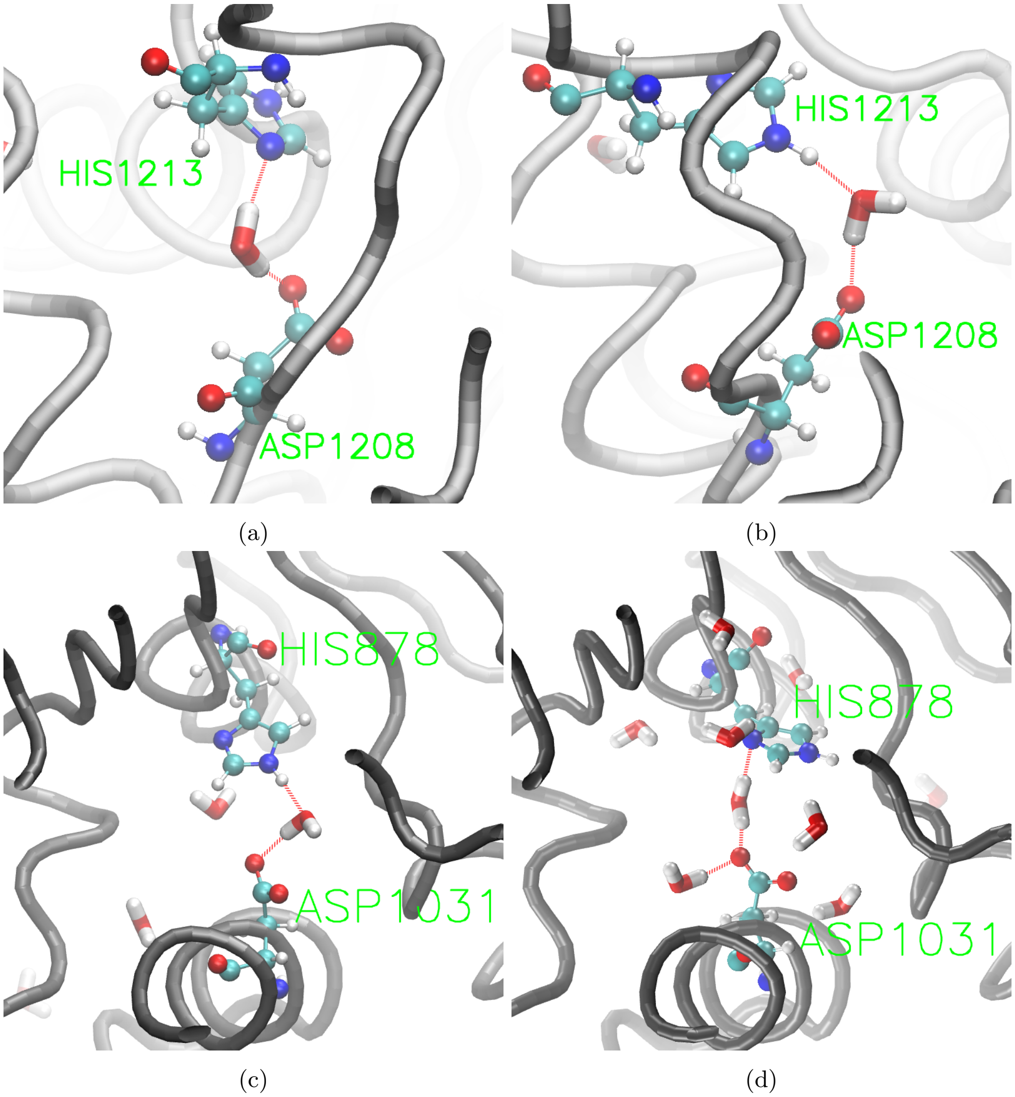


Figure S3: Water simultaneously forming hydrogen-bonds with (a,b) ASP1208 and HIS1213 in MtbFAS-I DH, and (c,d) HIS878 and ASP1031 in hFAS-I DH. Hydrogen bonds, shown as red lines, are identified using VMD with a cutoff distance of 3.0 Åand an angle of 20°.

Table S1: hFAS-I MAT and MtbFAS-I AT mapped residues table

| Sr. | hFAS-I | MtbFAS-I | QH | Sr. | hFAS-I | MtbFAS-I | QH | Sr. | hFAS-I | MtbFAS-I | QH | Sr. | hFAS-I | MtbFAS-I | QH |
|---|---|---|---|---|---|---|---|---|---|---|---|---|---|---|---|
| 1 | ARG468 | HIS1 | 0.51 | 58 | MET570 | GLN112 | 0.62 | 115 | VAL638 | PRO204 | 0.73 | 172 | GLU733 | ARG292 | 0.53 |
| 2 | ARG490 | GLU11 | 0.59 | 59 | GLY571 | GLY113 | 0.58 | 116 | VAL639 | VAL205 | 0.72 | 173 | TYR734 | GLU293 | 0.55 |
| 3 | PRO491 | PRO12 | 0.63 | 60 | PRO574 | PRO121 | 0.6 | 117 | PRO640 | LEU206 | 0.72 | 174 | ASN735 | LEU294 | 0.67 |
| 4 | LEU492 | TYR13 | 0.67 | 61 | ASP575 | VAL122 | 0.63 | 118 | ALA641 | SER207 | 0.72 | 175 | VAL736 | ALA295 | 0.55 |
| 5 | TRP493 | ALA14 | 0.72 | 62 | GLY576 | ALA123 | 0.66 | 119 | CYS642 | ILE208 | 0.68 | 176 | ASN737 | ASP296 | 0.56 |
| 6 | PHE494 | VAL15 | 0.72 | 63 | ILE577 | MET124 | 0.69 | 120 | HIS643 | ARG209 | 0.69 | 177 | ASN738 | ALA297 | 0.64 |
| 7 | ILE495 | ALA16 | 0.73 | 64 | VAL578 | ALA125 | 0.68 | 121 | ASN644 | ASN210 | 0.69 | 178 | LEU739 | LEU299 | 0.6 |
| 8 | CYS496 | PHE17 | 0.71 | 65 | GLY579 | GLY126 | 0.67 | 122 | SER645 | GLY211 | 0.67 | 179 | VAL740 | ILE300 | 0.62 |
| 9 | SER497 | GLY18 | 0.71 | 66 | HIS580 | HIS127 | 0.65 | 123 | LYS646 | ARG212 | 0.67 | 180 | SER741 | ARG301 | 0.55 |
| 10 | GLY498 | GLY19 | 0.62 | 67 | SER581 | SER128 | 0.61 | 124 | ASP647 | ARG213 | 0.66 | 181 | PRO742 | LYS302 | 0.58 |
| 11 | MET499 | GLN20 | 0.56 | 68 | LEU582 | GLN129 | 0.57 | 125 | THR648 | ALA214 | 0.61 | 182 | VAL743 | VAL303 | 0.64 |
| 12 | GLY500 | GLY21 | 0.68 | 69 | GLY583 | GLY130 | 0.71 | 126 | VAL649 | VAL215 | 0.63 | 183 | LEU744 | ASP304 | 0.64 |
| 13 | THR501 | SER22 | 0.67 | 70 | GLU584 | VAL131 | 0.68 | 127 | THR650 | VAL216 | 0.69 | 184 | PHE745 | TRP305 | 0.65 |
| 14 | GLN502 | ALA23 | 0.61 | 71 | VAL585 | LEU132 | 0.71 | 128 | ILE651 | ILE217 | 0.72 | 185 | GLN746 | VAL306 | 0.65 |
| 15 | GLY507 | LEU28 | 0.51 | 72 | ALA586 | ALA133 | 0.72 | 129 | SER652 | THR218 | 0.73 | 186 | GLU747 | ASP307 | 0.66 |
| 16 | LEU508 | GLU29 | 0.5 | 73 | CYS587 | VAL134 | 0.71 | 130 | GLY653 | GLY219 | 0.72 | 187 | ALA748 | GLU308 | 0.62 |
| 17 | MET511 | VAL32 | 0.54 | 74 | GLY588 | GLU135 | 0.7 | 131 | PRO654 | THR220 | 0.72 | 188 | LEU749 | ILE309 | 0.67 |
| 18 | LEU513 | THR35 | 0.54 | 75 | TYR589 | ALA136 | 0.7 | 132 | GLN655 | PRO221 | 0.71 | 189 | TRP750 | THR310 | 0.66 |
| 19 | ASP514 | GLY36 | 0.56 | 76 | ALA590 | LEU137 | 0.71 | 133 | ALA656 | GLU222 | 0.71 | 190 | HIS751 | ARG311 | 0.58 |
| 20 | PHE516 | ILE37 | 0.55 | 77 | ASP591 | LYS138 | 0.7 | 134 | PRO657 | GLN223 | 0.75 | 191 | VAL752 | VAL312 | 0.63 |
| 21 | ARG517 | GLU38 | 0.68 | 78 | LEU594 | ARG143 | 0.52 | 135 | VAL658 | LEU224 | 0.69 | 192 | PRO753 | GLY316 | 0.52 |
| 22 | ILE520 | LEU41 | 0.66 | 79 | GLN596 | ASP144 | 0.7 | 136 | PHE659 | SER225 | 0.56 | 193 | HIS755 | ARG318 | 0.52 |
| 23 | LEU521 | ALA42 | 0.7 | 80 | GLU597 | VAL145 | 0.74 | 137 | GLU660 | ARG226 | 0.67 | 194 | VAL757 | TRP319 | 0.72 |
| 24 | SER523 | LEU44 | 0.51 | 81 | GLU598 | GLU146 | 0.68 | 138 | PHE661 | PHE227 | 0.73 | 195 | VAL758 | ILE320 | 0.73 |
| 25 | ASP524 | VAL45 | 0.7 | 82 | ALA599 | LEU147 | 0.66 | 139 | VAL662 | GLU228 | 0.55 | 196 | LEU759 | LEU321 | 0.66 |
| 26 | GLU525 | GLY46 | 0.68 | 83 | VAL600 | PHE148 | 0.74 | 140 | GLN664 | TYR230 | 0.65 | 197 | GLU760 | ASP322 | 0.64 |
| 27 | VAL527 | ALA48 | 0.59 | 84 | LEU601 | ALA149 | 0.73 | 141 | LEU665 | CYS231 | 0.61 | 198 | ILE761 | LEU323 | 0.7 |
| 28 | LYS528 | GLU49 | 0.63 | 85 | ALA602 | LEU150 | 0.64 | 142 | GLU668 | ILE234 | 0.61 | 199 | ALA762 | GLY324 | 0.66 |
| 29 | LYS533 | GLU68 | 0.54 | 86 | ALA603 | ALA151 | 0.72 | 143 | PHE671 | VAL255 | 0.63 | 200 | PRO763 | PRO325 | 0.57 |
| 30 | VAL534 | PRO69 | 0.61 | 87 | TYR604 | GLN152 | 0.74 | 144 | ALA672 | PHE256 | 0.63 | 201 | HIS764 | GLY326 | 0.71 |
| 31 | SER535 | LEU70 | 0.68 | 88 | TRP605 | LEU153 | 0.73 | 145 | LYS673 | GLU257 | 0.6 | 202 | ALA765 | ASP327 | 0.64 |
| 32 | GLN536 | GLN71 | 0.68 | 89 | ARG606 | ILE154 | 0.73 | 146 | GLU674 | PRO258 | 0.59 | 203 | LEU766 | ILE328 | 0.68 |
| 33 | LEU537 | TRP72 | 0.51 | 90 | GLY607 | GLY155 | 0.7 | 147 | VAL675 | VAL259 | 0.59 | 204 | LEU767 | LEU329 | 0.59 |
| 34 | THR541 | VAL82 | 0.65 | 91 | GLN608 | ALA156 | 0.68 | 148 | ARG676 | GLN260 | 0.56 | 205 | GLN768 | THR330 | 0.63 |
| 35 | ASP542 | PRO83 | 0.59 | 92 | CYS609 | ALA157 | 0.7 | 149 | THR677 | VAL261 | 0.65 | 206 | ALA769 | ARG331 | 0.52 |
| 36 | PHE546 | LEU88 | 0.54 | 93 | ILE610 | GLY158 | 0.6 | 150 | GLY679 | GLU262 | 0.52 | 207 | VAL770 | LEU332 | 0.56 |
| 37 | ILE549 | ALA91 | 0.57 | 94 | LYS611 | THR159 | 0.52 | 151 | MET680 | VAL263 | 0.67 | 208 | LEU771 | THR333 | 0.62 |
| 38 | VAL550 | ALA92 | 0.68 | 95 | GLU612 | LEU160 | 0.54 | 152 | ALA681 | GLY264 | 0.69 | 209 | LYS772 | ALA334 | 0.65 |
| 39 | HIS551 | VAL93 | 0.63 | 96 | ALA613 | VAL161 | 0.56 | 153 | PHE682 | PHE265 | 0.67 | 210 | ARG773 | PRO335 | 0.58 |
| 40 | SER552 | SER94 | 0.61 | 97 | HIS614 | ARG163 | 0.64 | 154 | HIS683 | HIS266 | 0.71 | 211 | GLY774 | VAL336 | 0.58 |
| 41 | PHE553 | VAL95 | 0.65 | 98 | PRO616 | GLY166 | 0.7 | 155 | SER684 | THR267 | 0.73 | 212 | LEU775 | ILE337 | 0.65 |
| 42 | VAL554 | PRO96 | 0.62 | 99 | PRO617 | SER168 | 0.56 | 156 | TYR685 | PRO268 | 0.63 | 213 | LYS776 | LEU340 | 0.67 |
| 43 | SER555 | GLY97 | 0.61 | 100 | GLY618 | PRO175 | 0.62 | 157 | PHE686 | ARG269 | 0.71 | 214 | SER778 | GLY341 | 0.57 |
| 44 | LEU556 | VAL98 | 0.65 | 101 | MET620 | MET176 | 0.69 | 158 | MET687 | LEU270 | 0.67 | 215 | CYS779 | ILE342 | 0.67 |
| 45 | THR557 | LEU99 | 0.7 | 102 | ALA621 | VAL177 | 0.68 | 159 | GLU688 | SER271 | 0.63 | 216 | THR780 | GLY343 | 0.67 |
| 46 | ALA558 | LEU100 | 0.7 | 103 | ALA622 | SER178 | 0.62 | 160 | ALA689 | ASP272 | 0.58 | 217 | ILE781 | ILE344 | 0.68 |
| 47 | ILE559 | THR101 | 0.68 | 104 | VAL623 | VAL179 | 0.6 | 161 | ILE690 | GLY273 | 0.61 | 218 | ILE782 | VAL345 | 0.74 |
| 48 | GLN560 | GLN102 | 0.73 | 105 | LEU625 | ALA182 | 0.53 | 162 | ALA691 | ILE274 | 0.6 | 219 | PRO783 | PRO346 | 0.7 |
| 49 | ILE561 | ILE103 | 0.74 | 106 | SER626 | ASP183 | 0.6 | 163 | PRO692 | ASP275 | 0.65 | 220 | LEU784 | ALA347 | 0.65 |
| 50 | GLY562 | ALA104 | 0.72 | 107 | TRP627 | PRO184 | 0.6 | 164 | PRO693 | ILE276 | 0.65 | 221 | GLY799 | GLY352 | 0.6 |
| 51 | LEU563 | ALA105 | 0.71 | 108 | GLU628 | GLU185 | 0.59 | 165 | LEU694 | VAL277 | 0.63 | 222 | ILE800 | GLN353 | 0.64 |
| 52 | ILE564 | THR106 | 0.73 | 109 | GLU629 | ARG186 | 0.62 | 166 | LEU695 | ALA278 | 0.53 | 223 | GLY801 | ARG354 | 0.59 |
| 53 | ASP565 | ARG107 | 0.71 | 110 | CYS630 | ILE187 | 0.62 | 167 | GLN696 | GLY279 | 0.57 | 224 | ARG802 | ASN355 | 0.66 |
| 54 | LEU566 | ALA108 | 0.69 | 111 | LYS631 | GLY188 | 0.62 | 168 | GLU697 | TRP280 | 0.56 | 225 | LEU803 | LEU356 | 0.64 |
| 55 | LEU567 | LEU109 | 0.7 | 112 | GLN632 | ARG189 | 0.62 | 169 | VAL701 | LEU286 | 0.54 | 226 | HIS804 | PHE357 | 0.6 |
| 56 | SER568 | ALA110 | 0.65 | 113 | ARG633 | LEU190 | 0.63 | 170 | SER731 | LEU290 | 0.57 | | | | |
| 57 | CYS569 | ARG111 | 0.67 | 114 | CYS634 | LEU191 | 0.64 | 171 | ALA732 | ALA291 | 0.59 | | | | |

### Table S2: hFAS-I MAT and MtbFAS-I MPT mapped residues table

| S.N | hFAS-I | MtbFAS-I | QH | S.N | hFAS-I | MtbFAS-I | QH | S.N | hFAS-I | MtbFAS-I | QH | S.N | hFAS-I | MtbFAS-I | QH |
|---|---|---|---|---|---|---|---|---|---|---|---|---|---|---|---|
| 1 | SER730 | PHE1628 | 0.5 | 58 | MET511 | ARG1330 | 0.6 | 115 | VAL701 | VAL1563 | 0.67 | 172 | PHE516 | ALA1336 | 0.71 |
| 2 | GLY571 | GLY1405 | 0.5 | 59 | SER544 | GLY1378 | 0.6 | 116 | GLN746 | ILE1643 | 0.67 | 173 | ASP518 | LYS1338 | 0.71 |
| 3 | GLN768 | ALA1675 | 0.5 | 60 | ARG708 | ILE1573 | 0.61 | 117 | ARG802 | VAL1703 | 0.67 | 174 | ARG515 | ALA1335 | 0.71 |
| 4 | TRP722 | ARG1584 | 0.51 | 61 | VAL639 | GLU1494 | 0.61 | 118 | PHE546 | LEU1380 | 0.67 | 175 | LEU539 | ARG1360 | 0.71 |
| 5 | MET785 | GLU1698 | 0.51 | 62 | SER714 | PRO1579 | 0.61 | 119 | LEU698 | LEU1560 | 0.68 | 176 | VAL554 | VAL1388 | 0.71 |
| 6 | PRO616 | PRO1452 | 0.51 | 63 | GLY801 | ALA1702 | 0.61 | 120 | ILE782 | LEU1695 | 0.68 | 177 | ALA748 | THR1645 | 0.71 |
| 7 | SER568 | ARG1402 | 0.52 | 64 | LEU508 | MET1327 | 0.61 | 121 | LYS700 | ARG1562 | 0.68 | 178 | THR780 | GLU1693 | 0.71 |
| 8 | ALA762 | GLY1667 | 0.52 | 65 | VAL770 | LEU1677 | 0.61 | 122 | VAL585 | TYR1421 | 0.68 | 179 | SER552 | THR1386 | 0.71 |
| 9 | PRO753 | GLY1659 | 0.52 | 66 | LEU563 | GLN1397 | 0.62 | 123 | THR557 | ALA1391 | 0.68 | 180 | GLU747 | GLU1644 | 0.71 |
| 10 | MET620 | LEU1463 | 0.52 | 67 | ILE561 | ALA1395 | 0.62 | 124 | ALA558 | THR1392 | 0.68 | 181 | ASP548 | LEU1382 | 0.71 |
| 11 | ALA526 | PHE1346 | 0.53 | 68 | LEU521 | ASP1341 | 0.62 | 125 | TYR734 | LEU1631 | 0.68 | 182 | GLN536 | HIS1357 | 0.71 |
| 12 | MET680 | VAL1542 | 0.53 | 69 | SER581 | SER1417 | 0.62 | 126 | LYS699 | ASP1561 | 0.68 | 183 | LYS533 | SER1354 | 0.72 |
| 13 | GLY774 | THR1681 | 0.53 | 70 | SER523 | ALA1343 | 0.62 | 127 | HIS751 | LEU1648 | 0.68 | 184 | HIS683 | HIS1545 | 0.72 |
| 14 | ILE577 | ALA1413 | 0.54 | 71 | LEU513 | SER1333 | 0.62 | 128 | GLY799 | ASP1700 | 0.68 | 185 | VAL534 | VAL1355 | 0.72 |
| 15 | SER652 | ALA1508 | 0.54 | 72 | LYS772 | THR1679 | 0.62 | 129 | LEU771 | ALA1678 | 0.68 | 186 | CYS642 | ASN1497 | 0.72 |
| 16 | MET570 | GLN1404 | 0.55 | 73 | ALA681 | PRO1543 | 0.62 | 130 | SER778 | THR1691 | 0.68 | 187 | MET687 | LEU1549 | 0.72 |
| 17 | LYS673 | ILE1535 | 0.55 | 74 | LEU582 | VAL1418 | 0.63 | 131 | ASP514 | LYS1334 | 0.68 | 188 | TRP605 | HIS1442 | 0.73 |
| 18 | CYS569 | GLU1403 | 0.55 | 75 | LEU510 | VAL1329 | 0.63 | 132 | THR648 | GLN1504 | 0.68 | 189 | SER519 | VAL1339 | 0.73 |
| 19 | ALA621 | ALA1464 | 0.55 | 76 | MET499 | GLN1318 | 0.63 | 133 | LEU766 | THR1673 | 0.68 | 190 | GLN696 | ARG1558 | 0.73 |
| 20 | THR715 | ASN1580 | 0.56 | 77 | CYS587 | ALA1423 | 0.63 | 134 | LEU759 | VAL1664 | 0.68 | 191 | ASN644 | ASN1499 | 0.73 |
| 21 | HIS804 | ALA1706 | 0.56 | 78 | GLN560 | ALA1394 | 0.63 | 135 | THR541 | ASN1362 | 0.68 | 192 | LEU695 | ARG1557 | 0.73 |
| 22 | ILE564 | VAL1398 | 0.56 | 79 | TRP712 | TYR1577 | 0.63 | 136 | CYS496 | PHE1315 | 0.69 | 193 | GLY498 | GLY1317 | 0.73 |
| 23 | ARG711 | ARG1576 | 0.56 | 80 | GLY562 | ALA1396 | 0.63 | 137 | SER595 | GLN1432 | 0.69 | 194 | ALA641 | VAL1496 | 0.73 |
| 24 | GLU525 | LYS1345 | 0.56 | 81 | GLU733 | GLU1630 | 0.63 | 138 | GLU597 | GLU1434 | 0.69 | 195 | HIS643 | PHE1498 | 0.73 |
| 25 | VAL527 | THR1347 | 0.56 | 82 | GLY637 | PHE1492 | 0.63 | 139 | GLY592 | GLY1429 | 0.69 | 196 | TYR685 | ARG1547 | 0.73 |
| 26 | VAL638 | LEU1493 | 0.56 | 83 | ILE520 | TRP1340 | 0.64 | 140 | ILE559 | VAL1393 | 0.69 | 197 | GLN608 | SER1445 | 0.74 |
| 27 | ARG703 | PRO1565 | 0.56 | 84 | ASP547 | TYR1381 | 0.64 | 141 | VAL752 | LEU1649 | 0.69 | 198 | LEU694 | PHE1556 | 0.74 |
| 28 | LYS528 | ARG1348 | 0.56 | 85 | ASP591 | THR1428 | 0.64 | 142 | VAL736 | ALA1633 | 0.69 | 199 | PRO640 | ILE1495 | 0.74 |
| 29 | LEU594 | TYR1431 | 0.57 | 86 | LEU784 | ALA1697 | 0.64 | 143 | THR650 | ALA1506 | 0.69 | 200 | ASN737 | TRP1634 | 0.74 |
| 30 | ARG773 | ASN1680 | 0.57 | 87 | SER716 | LEU1581 | 0.64 | 144 | ILE781 | VAL1694 | 0.69 | 201 | CYS609 | LYS1446 | 0.74 |
| 31 | VAL578 | CYS1414 | 0.57 | 88 | GLY507 | GLY1326 | 0.64 | 145 | LEU537 | VAL1358 | 0.69 | 202 | VAL550 | GLN1384 | 0.74 |
| 32 | GLY576 | ILE1412 | 0.57 | 89 | GLY500 | GLY1319 | 0.64 | 146 | LEU538 | VAL1359 | 0.69 | 203 | LEU744 | ARG1641 | 0.74 |
| 33 | ALA590 | VAL1427 | 0.57 | 90 | GLU598 | ALA1435 | 0.65 | 147 | VAL623 | ILE1466 | 0.69 | 204 | HIS551 | PHE1385 | 0.74 |
| 34 | LEU566 | GLU1400 | 0.57 | 91 | GLY505 | GLY1324 | 0.65 | 148 | PHE530 | LEU1351 | 0.69 | 205 | PRO693 | GLU1555 | 0.74 |
| 35 | GLU612 | ASP1449 | 0.57 | 92 | PHE671 | SER1533 | 0.65 | 149 | LYS646 | ARG1501 | 0.69 | 206 | PRO692 | ALA1554 | 0.74 |
| 36 | ALA613 | ILE1450 | 0.57 | 93 | ALA586 | THR1422 | 0.65 | 150 | ALA603 | VAL1440 | 0.7 | 207 | THR545 | VAL1379 | 0.74 |
| 37 | GLU660 | GLU1515 | 0.57 | 94 | LEU601 | GLU1438 | 0.65 | 151 | VAL758 | PHE1663 | 0.7 | 208 | GLY624 | ARG1467 | 0.74 |
| 38 | ALA672 | PHE1534 | 0.57 | 95 | LEU713 | ILE1578 | 0.65 | 152 | SER497 | PRO1316 | 0.7 | 209 | ARG606 | ARG1443 | 0.75 |
| 39 | GLU668 | ARG1524 | 0.57 | 96 | HIS580 | HIS1416 | 0.65 | 153 | SER535 | LEU1356 | 0.7 | 210 | VAL743 | VAL1640 | 0.75 |
| 40 | ASP565 | ALA1399 | 0.58 | 97 | ILE651 | ILE1507 | 0.65 | 154 | SER645 | LEU1500 | 0.7 | 211 | ILE610 | MET1447 | 0.75 |
| 41 | LEU567 | MET1401 | 0.58 | 98 | ASN735 | LEU1632 | 0.65 | 155 | GLU584 | GLU1420 | 0.7 | 212 | ASN738 | GLN1635 | 0.75 |
| 42 | ASP542 | PRO1363 | 0.58 | 99 | CYS593 | ILE1430 | 0.65 | 156 | LEU556 | MET1390 | 0.7 | 213 | VAL649 | TYR1505 | 0.75 |
| 43 | PRO529 | THR1350 | 0.58 | 100 | SER724 | PHE1592 | 0.65 | 157 | PHE553 | GLN1387 | 0.7 | 214 | ALA689 | VAL1551 | 0.75 |
| 44 | GLN502 | GLN1321 | 0.59 | 101 | ASP647 | GLY1502 | 0.65 | 158 | VAL757 | ARG1662 | 0.7 | 215 | GLY531 | GLY1352 | 0.75 |
| 45 | TRP503 | HIS1322 | 0.59 | 102 | LEU532 | PHE1353 | 0.65 | 159 | LEU749 | GLN1646 | 0.7 | 216 | GLY607 | GLY1444 | 0.76 |
| 46 | ALA622 | ALA1465 | 0.59 | 103 | ARG522 | THR1342 | 0.66 | 160 | PHE745 | TRP1642 | 0.7 | 217 | LEU615 | VAL1451 | 0.76 |
| 47 | THR501 | ILE1320 | 0.59 | 104 | ARG512 | ALA1331 | 0.66 | 161 | LYS611 | HIS1448 | 0.7 | 218 | ALA691 | VAL1553 | 0.76 |
| 48 | LEU767 | VAL1674 | 0.59 | 105 | VAL600 | LEU1437 | 0.66 | 162 | CYS779 | VAL1692 | 0.7 | 219 | VAL740 | ALA1637 | 0.76 |
| 49 | ALA765 | PRO1672 | 0.59 | 106 | ALA599 | LEU1436 | 0.66 | 163 | TRP750 | ASP1647 | 0.7 | 220 | ILE690 | GLY1552 | 0.76 |
| 50 | SER540 | ASP1361 | 0.59 | 107 | GLY583 | GLY1419 | 0.66 | 164 | MET506 | MET1325 | 0.7 | 221 | SER684 | SER1546 | 0.76 |
| 51 | ASP524 | ASP1344 | 0.6 | 108 | PHE686 | VAL1548 | 0.66 | 165 | GLU760 | GLU1665 | 0.7 | 222 | LEU739 | PHE1636 | 0.77 |
| 52 | ILE702 | MET1564 | 0.6 | 109 | ALA602 | MET1439 | 0.67 | 166 | ILE549 | THR1383 | 0.7 | 223 | SER741 | SER1638 | 0.77 |
| 53 | ARG504 | LYS1323 | 0.6 | 110 | ARG517 | ARG1337 | 0.67 | 167 | PHE682 | PHE1544 | 0.7 | 224 | GLU688 | ARG1550 | 0.77 |
| 54 | SER509 | GLU1328 | 0.6 | 111 | TYR589 | ALA1425 | 0.67 | 168 | TYR604 | PHE1441 | 0.71 | 225 | PRO742 | PRO1639 | 0.78 |
| 55 | ALA710 | GLY1575 | 0.6 | 112 | GLY588 | LEU1424 | 0.67 | 169 | GLN596 | LEU1433 | 0.71 | | | | |
| 56 | ILE800 | ALA1701 | 0.6 | 113 | ILE761 | ILE1666 | 0.67 | 170 | SER555 | ALA1389 | 0.71 | | | | |
| 57 | PRO783 | ASN1696 | 0.6 | 114 | GLY579 | GLY1415 | 0.67 | 171 | GLU697 | SER1559 | 0.71 | | | | |

### Table S3: hFAS-I KS and MtbFAS-I KS mapped residues table

| Sr. | HFAS-I | MtbFAS-I | QH | Sr. | HFAS-I | MtbFAS-I | QH | Sr. | HFAS-I | MtbFAS-I | QH | Sr. | HFAS-I | MtbFAS-I | QH |
|---|---|---|---|---|---|---|---|---|---|---|---|---|---|---|---|
| 1 | MET1 | ASP2418 | 0.61 | 101 | ARG143 | HIS2680 | 0.66 | 201 | ALA249 | ALA2801 | 0.7 | 301 | VAL377 | VAL2936 | 0.68 |
| 2 | GLU3 | LEU2419 | 0.81 | 102 | LEU144 | VAL2681 | 0.67 | 202 | GLY250 | GLN2802 | 0.73 | 302 | ASP378 | ARG2937 | 0.72 |
| 3 | VAL4 | VAL2420 | 0.81 | 103 | SER145 | VAL2682 | 0.66 | 203 | THR251 | SER2803 | 0.73 | 303 | GLN379 | ASP2938 | 0.66 |
| 4 | VAL5 | VAL2421 | 0.82 | 104 | PHE146 | GLN2683 | 0.7 | 204 | ASN252 | PHE2804 | 0.77 | 304 | PRO380 | THR2939 | 0.68 |
| 5 | ILE6 | ILE2422 | 0.83 | 105 | PHE147 | SER2684 | 0.67 | 205 | THR253 | GLY2805 | 0.72 | 305 | LEU381 | LEU2940 | 0.7 |
| 6 | ALA7 | VAL2423 | 0.83 | 106 | ASP149 | GLY2687 | 0.56 | 206 | ASP254 | ASP2806 | 0.73 | 306 | PRO382 | ARG2941 | 0.62 |
| 7 | GLY8 | GLY2424 | 0.79 | 107 | PHE150 | SER2688 | 0.73 | 207 | GLY255 | GLY2807 | 0.69 | 307 | GLY385 | LEU2948 | 0.76 |
| 8 | MET9 | GLY2425 | 0.78 | 108 | ARG151 | TYR2689 | 0.78 | 208 | PHE256 | VAL2808 | 0.68 | 308 | GLY386 | LYS2949 | 0.73 |
| 9 | SER10 | ALA2426 | 0.74 | 109 | GLY152 | GLY2690 | 0.75 | 209 | LYS257 | THR2810 | 0.73 | 309 | ASN387 | ALA2950 | 0.77 |
| 10 | GLY11 | GLU2427 | 0.75 | 110 | PRO153 | ALA2691 | 0.74 | 210 | GLY266 | PRO2815 | 0.61 | 310 | VAL388 | GLY2951 | 0.82 |
| 11 | LYS12 | ILE2428 | 0.74 | 111 | SER154 | MET2692 | 0.73 | 211 | GLU270 | LEU2820 | 0.5 | 311 | GLY389 | MET2952 | 0.82 |
| 12 | LEU13 | GLY2429 | 0.75 | 112 | ILE155 | ILE2693 | 0.67 | 212 | GLN271 | GLY2821 | 0.56 | 312 | ILE390 | LEU2953 | 0.8 |
| 13 | PRO14 | PRO2430 | 0.77 | 113 | ALA156 | HIS2694 | 0.69 | 213 | LEU276 | LEU2831 | 0.63 | 313 | ASN391 | THR2954 | 0.78 |
| 14 | SER16 | TYR2431 | 0.65 | 114 | LEU157 | PRO2695 | 0.71 | 214 | GLN278 | ALA2832 | 0.49 | 314 | SER392 | SER2955 | 0.77 |
| 15 | GLU17 | GLY2432 | 0.53 | 115 | ASP158 | VAL2696 | 0.78 | 215 | ALA280 | ARG2833 | 0.65 | 315 | PHE393 | LEU2956 | 0.8 |
| 16 | ASN18 | SER2433 | 0.69 | 116 | THR159 | ALA2697 | 0.76 | 216 | VAL282 | LEU2835 | 0.57 | 316 | GLY394 | GLY2957 | 0.74 |
| 17 | LEU19 | SER2434 | 0.73 | 117 | ALA160 | ALA2698 | 0.73 | 217 | ALA283 | ALA2841 | 0.69 | 317 | PHE395 | PHE2958 | 0.5 |
| 18 | GLN20 | ARG2435 | 0.72 | 118 | CYS161 | CYS2699 | 0.77 | 218 | PRO284 | ALA2842 | 0.74 | 318 | GLY396 | GLY2959 | 0.57 |
| 19 | GLU21 | THR2436 | 0.74 | 119 | SER162 | ALA2700 | 0.8 | 219 | GLU285 | ASP2843 | 0.72 | 319 | GLY397 | HIS2960 | 0.63 |
| 20 | PHE22 | ARG2437 | 0.77 | 120 | SER163 | THR2701 | 0.78 | 220 | SER286 | ASP2844 | 0.79 | 320 | SER398 | VAL2961 | 0.74 |
| 21 | TRP23 | PHE2438 | 0.74 | 121 | SER164 | ALA2702 | 0.75 | 221 | PHE287 | VAL2845 | 0.76 | 321 | ASN399 | SER2962 | 0.78 |
| 22 | ASP24 | GLU2439 | 0.72 | 122 | LEU165 | ALA2703 | 0.77 | 222 | GLU288 | ALA2846 | 0.8 | 322 | VAL400 | GLY2963 | 0.78 |
| 23 | ASN25 | MET2440 | 0.74 | 123 | MET166 | VAL2704 | 0.79 | 223 | TYR289 | VAL2847 | 0.77 | 323 | HIS401 | LEU2964 | 0.81 |
| 24 | LEU26 | GLU2441 | 0.75 | 124 | ALA167 | SER2705 | 0.8 | 224 | ILE290 | ILE2848 | 0.81 | 324 | ILE402 | VAL2965 | 0.81 |
| 25 | ILE27 | VAL2442 | 0.73 | 125 | LEU168 | VAL2706 | 0.79 | 225 | GLU291 | SER2849 | 0.8 | 325 | ILE403 | ALA2966 | 0.8 |
| 26 | GLY28 | GLU2443 | 0.61 | 126 | GLN169 | GLU2707 | 0.78 | 226 | ALA292 | LYS2850 | 0.78 | 326 | LEU404 | LEU2967 | 0.78 |
| 27 | GLY29 | ASN2444 | 0.54 | 127 | ASN170 | GLU2708 | 0.79 | 227 | HIS293 | HIS2851 | 0.77 | 327 | ARG405 | VAL2968 | 0.79 |
| 28 | MET32 | GLY2498 | 0.63 | 128 | ALA171 | GLY2709 | 0.79 | 228 | GLY294 | ASP2852 | 0.67 | 328 | PRO406 | HIS2969 | 0.82 |
| 29 | VAL33 | ILE2499 | 0.74 | 129 | TYR172 | VAL2710 | 0.76 | 229 | THR295 | THR2853 | 0.68 | | | | |
| 30 | THR34 | ARG2500 | 0.75 | 130 | GLN173 | ASP2711 | 0.76 | 230 | GLY296 | SER2854 | 0.71 | | | | |
| 31 | ASP35 | GLU2501 | 0.7 | 131 | ALA174 | LYS2712 | 0.78 | 231 | THR297 | THR2855 | 0.73 | | | | |
| 32 | ARG38 | VAL2503 | 0.76 | 132 | ILE175 | ILE2713 | 0.77 | 232 | LYS298 | LEU2856 | 0.74 | | | | |
| 33 | TYR45 | LEU2576 | 0.65 | 133 | HIS176 | ARG2714 | 0.75 | 233 | VAL299 | ALA2857 | 0.73 | | | | |
| 34 | GLY46 | SER2577 | 0.53 | 134 | SER177 | LEU2715 | 0.73 | 234 | GLY300 | ASN2858 | 0.74 | | | | |
| 35 | ARG50 | VAL2580 | 0.71 | 135 | GLY178 | GLY2716 | 0.77 | 235 | ASP301 | ASP2859 | 0.75 | | | | |
| 36 | SER51 | GLY2581 | 0.75 | 136 | GLN179 | LYS2717 | 0.77 | 236 | PRO302 | PRO2860 | 0.77 | | | | |
| 37 | GLY52 | GLY2582 | 0.76 | 137 | CYS180 | ALA2718 | 0.79 | 237 | GLN303 | ASN2861 | 0.77 | | | | |
| 38 | LYS53 | GLN2583 | 0.77 | 138 | PRO181 | GLN2719 | 0.78 | 238 | GLU304 | GLU2862 | 0.77 | | | | |
| 39 | LEU54 | ILE2584 | 0.76 | 139 | ALA182 | LEU2720 | 0.8 | 239 | LEU305 | THR2863 | 0.79 | | | | |
| 40 | LEU57 | PHE2588 | 0.49 | 140 | ALA183 | VAL2721 | 0.8 | 240 | ASN306 | GLU2864 | 0.78 | | | | |
| 41 | ARG59 | ASP2589 | 0.61 | 141 | ILE184 | VAL2722 | 0.81 | 241 | GLY307 | LEU2865 | 0.7 | | | | |
| 42 | PHE60 | PRO2590 | 0.56 | 142 | VAL185 | ALA2723 | 0.79 | 242 | ILE308 | HIS2866 | 0.68 | | | | |
| 43 | ASP61 | GLY2594 | 0.59 | 143 | GLY186 | GLY2724 | 0.62 | 243 | THR309 | GLU2867 | 0.69 | | | | |
| 44 | HIS73 | GLY2601 | 0.69 | 144 | GLY187 | GLY2725 | 0.78 | 244 | ARG310 | ARG2868 | 0.54 | | | | |
| 45 | THR74 | SER2602 | 0.64 | 145 | ILE188 | LEU2726 | 0.77 | 245 | LEU312 | ALA2870 | 0.74 | | | | |
| 46 | MET75 | ILE2603 | 0.72 | 146 | ASN189 | ASP2727 | 0.76 | 246 | CYS313 | ARG2875 | 0.52 | | | | |
| 47 | ASP76 | ASP2604 | 0.7 | 147 | VAL190 | ASP2728 | 0.74 | 247 | THR315 | SER2876 | 0.73 | | | | |
| 48 | PRO77 | ARG2605 | 0.76 | 148 | LEU191 | LEU2729 | 0.65 | 248 | PRO319 | PRO2880 | 0.74 | | | | |
| 49 | GLN78 | LEU2606 | 0.78 | 149 | PRO194 | LEU2731 | 0.69 | 249 | LEU320 | LEU2881 | 0.71 | | | | |
| 50 | LEU79 | ALA2607 | 0.78 | 150 | ASN195 | GLU2732 | 0.7 | 250 | LEU321 | PHE2882 | 0.72 | | | | |
| 51 | ARG80 | VAL2608 | 0.73 | 151 | THR196 | GLY2733 | 0.64 | 251 | ILE322 | VAL2883 | 0.76 | | | | |
| 52 | LEU81 | TRP2609 | 0.74 | 152 | SER197 | ILE2734 | 0.55 | 252 | GLY323 | VAL2884 | 0.61 | | | | |
| 53 | LEU82 | ASN2610 | 0.76 | 153 | VAL198 | ILE2735 | 0.6 | 253 | SER324 | SER2885 | 0.74 | | | | |
| 54 | LEU83 | MET2611 | 0.72 | 154 | GLN199 | GLY2736 | 0.64 | 254 | THR325 | GLN2886 | 0.77 | | | | |
| 55 | GLU84 | VAL2612 | 0.67 | 155 | PHE200 | PHE2737 | 0.56 | 255 | LYS326 | LYS2887 | 0.78 | | | | |
| 56 | VAL85 | ALA2613 | 0.72 | 156 | LEU201 | GLY2738 | 0.65 | 256 | SER327 | SER2888 | 0.76 | | | | |
| 57 | THR86 | THR2614 | 0.77 | 157 | ARG202 | ASP2739 | 0.63 | 257 | ASN328 | LEU2889 | 0.74 | | | | |
| 58 | TYR87 | VAL2615 | 0.69 | 158 | LEU203 | MET2740 | 0.59 | 258 | MET329 | THR2890 | 0.74 | | | | |
| 59 | GLU88 | ASP2616 | 0.67 | 159 | GLY204 | ALA2741 | 0.69 | 259 | GLY330 | GLY2891 | 0.77 | | | | |
| 60 | ALA89 | ALA2617 | 0.76 | 160 | MET205 | ALA2742 | 0.76 | 260 | HIS331 | HIS2892 | 0.79 | | | | |
| 61 | ILE90 | PHE2618 | 0.73 | 161 | LEU206 | THR2743 | 0.77 | 261 | PRO332 | ALA2893 | 0.8 | | | | |
| 62 | VAL91 | LEU2619 | 0.69 | 162 | SER207 | ALA2744 | 0.73 | 262 | GLU333 | LYS2894 | 0.75 | | | | |
| 63 | ASP92 | SER2620 | 0.68 | 163 | PRO208 | ASP2745 | 0.72 | 263 | PRO334 | GLY2895 | 0.73 | | | | |
| 64 | GLY93 | SER2621 | 0.65 | 164 | THR211 | PHE2759 | 0.69 | 264 | ALA335 | GLY2896 | 0.79 | | | | |
| 65 | GLY94 | GLY2622 | 0.69 | 165 | CYS212 | SER2760 | 0.73 | 265 | SER336 | ALA2897 | 0.79 | | | | |
| 66 | ILE95 | PHE2623 | 0.69 | 166 | LYS213 | ARG2761 | 0.66 | 266 | GLY337 | ALA2898 | 0.78 | | | | |
| 67 | SER99 | GLU2627 | 0.62 | 167 | ALA214 | PRO2762 | 0.76 | 267 | LEU338 | VAL2899 | 0.77 | | | | |
| 68 | LEU100 | VAL2628 | 0.6 | 168 | PHE215 | ASN2763 | 0.75 | 268 | ALA339 | PHE2900 | 0.78 | | | | |
| 69 | ARG101 | MET2629 | 0.57 | 169 | ASP216 | ASP2764 | 0.76 | 269 | ALA340 | GLN2901 | 0.76 | | | | |
| 70 | THR103 | PRO2634 | 0.7 | 170 | THR217 | ARG2765 | 0.75 | 270 | LEU341 | MET2902 | 0.76 | | | | |
| 71 | THR105 | VAL2637 | 0.75 | 171 | ALA218 | ARG2766 | 0.67 | 271 | ALA342 | MET2903 | 0.79 | | | | |
| 72 | GLY106 | ALA2638 | 0.76 | 172 | GLY219 | ARG2767 | 0.65 | 272 | LYS343 | GLY2904 | 0.77 | | | | |
| 73 | VAL107 | ASN2639 | 0.77 | 173 | ASN220 | LEU2768 | 0.74 | 273 | VAL344 | LEU2905 | 0.77 | | | | |
| 74 | TRP108 | THR2640 | 0.71 | 174 | GLY221 | GLY2769 | 0.75 | 274 | LEU345 | CYS2906 | 0.77 | | | | |
| 75 | VAL109 | GLN2641 | 0.68 | 175 | TYR222 | PHE2770 | 0.77 | 275 | LEU346 | GLN2907 | 0.75 | | | | |
| 76 | GLY110 | GLY2642 | 0.75 | 176 | CYS223 | VAL2771 | 0.76 | 276 | SER347 | ILE2908 | 0.75 | | | | |
| 77 | VAL111 | THR2643 | 0.77 | 177 | ARG224 | GLU2772 | 0.77 | 277 | LEU348 | LEU2909 | 0.73 | | | | |
| 78 | SER112 | GLY2644 | 0.71 | 178 | SER225 | ALA2773 | 0.77 | 278 | GLU349 | ARG2910 | 0.72 | | | | |
| 79 | GLY113 | MET2645 | 0.68 | 179 | GLU226 | GLN2774 | 0.77 | 279 | HIS350 | ASP2911 | 0.73 | | | | |
| 80 | SER114 | GLY2647 | 0.59 | 180 | GLY227 | GLY2775 | 0.71 | 280 | GLY351 | GLY2912 | 0.74 | | | | |
| 81 | GLU115 | GLY2648 | 0.61 | 181 | VAL228 | GLY2776 | 0.59 | 281 | LEU352 | VAL2913 | 0.8 | | | | |
| 82 | THR116 | THR2649 | 0.55 | 182 | VAL229 | GLY2777 | 0.79 | 282 | TRP353 | ILE2914 | 0.75 | | | | |
| 83 | SER117 | SER2650 | 0.72 | 183 | ALA230 | THR2778 | 0.81 | 283 | ALA354 | PRO2915 | 0.74 | | | | |
| 84 | GLU118 | MET2651 | 0.61 | 184 | VAL231 | ILE2779 | 0.79 | 284 | PRO355 | PRO2916 | 0.71 | | | | |
| 85 | SER121 | MET2654 | 0.53 | 185 | LEU232 | LEU2780 | 0.79 | 285 | ASN356 | ASN2917 | 0.72 | | | | |
| 86 | PRO124 | ASN2663 | 0.57 | 186 | LEU233 | LEU2781 | 0.8 | 286 | LEU357 | ARG2918 | 0.63 | | | | |
| 87 | LEU127 | LYS2664 | 0.66 | 187 | THR234 | ALA2782 | 0.8 | 287 | HIS358 | SER2919 | 0.64 | | | | |
| 88 | VAL128 | PRO2665 | 0.63 | 188 | LYS235 | ARG2783 | 0.79 | 288 | PHE359 | LEU2920 | 0.68 | | | | |
| 89 | GLY129 | ASN2666 | 0.59 | 189 | LYS236 | GLY2784 | 0.79 | 289 | HIS360 | ASP2921 | 0.61 | | | | |
| 90 | TYR130 | ASP2667 | 0.59 | 190 | SER237 | ASP2785 | 0.76 | 290 | SER361 | CYS2922 | 0.63 | | | | |
| 91 | SER131 | ILE2668 | 0.64 | 191 | LEU238 | LEU2786 | 0.75 | 291 | PRO362 | VAL2923 | 0.68 | | | | |
| 92 | GLY134 | GLU2671 | 0.5 | 192 | ALA239 | ALA2787 | 0.7 | 292 | ASN363 | ASP2924 | 0.74 | | | | |
| 93 | CYS135 | VAL2672 | 0.5 | 193 | ARG241 | PRO2793 | 0.77 | 293 | PRO364 | ASP2925 | 0.76 | | | | |
| 94 | GLN136 | LEU2673 | 0.69 | 194 | VAL242 | VAL2794 | 0.8 | 294 | GLU365 | GLU2926 | 0.71 | | | | |
| 95 | ARG137 | PRO2674 | 0.74 | 195 | TYR243 | LEU2795 | 0.78 | 295 | ILE366 | LEU2927 | 0.53 | | | | |
| 96 | ALA138 | ASN2675 | 0.69 | 196 | ALA244 | ALA2796 | 0.78 | 296 | PRO367 | GLY2929 | 0.74 | | | | |
| 97 | MET139 | ILE2676 | 0.61 | 197 | THR245 | VAL2797 | 0.79 | 297 | ALA368 | SER2930 | 0.77 | | | | |
| 98 | MET140 | ILE2677 | 0.59 | 198 | ILE246 | VAL2798 | 0.79 | 298 | LEU374 | PHE2933 | 0.71 | | | | |
| 99 | ALA141 | ALA2678 | 0.58 | 199 | LEU247 | ALA2799 | 0.75 | 299 | GLN375 | VAL2934 | 0.77 | | | | |
| 100 | ASN142 | ALA2679 | 0.59 | 200 | ASN248 | PHE2800 | 0.71 | 300 | VAL376 | TRP2935 | 0.78 | | | | |

### Table S4: hFAS-I ACP and MtbFAS-I ACP mapped residues table

| Sr. | HFAS-I | MtbFAS-I | QH | Sr. | HFAS-I | MtbFAS-I | QH | Sr. | HFAS-I | MtbFAS-I | QH | Sr. | HFAS-I | MtbFAS-I | QH |
|---|---|---|---|---|---|---|---|---|---|---|---|---|---|---|---|
| 1 | GLN2124 | ALA1757 | 0.58 | 7 | ARG2138 | ARG1769 | 0.63 | 13 | SER2148 | SER1778 | 0.59 | 19 | LEU2187 | GLN1820 | 0.53 |
| 2 | VAL2128 | LEU1759 | 0.53 | 8 | LEU2140 | ILE1770 | 0.56 | 14 | THR2165 | GLU1800 | 0.51 | 20 | GLU2189 | THR1822 | 0.57 |
| 3 | GLU2129 | ALA1760 | 0.52 | 9 | ALA2141 | ASP1771 | 0.5 | 15 | THR2183 | GLY1816 | 0.53 | 21 | LEU2190 | LYS1823 | 0.71 |
| 4 | LEU2135 | ALA1766 | 0.5 | 10 | LEU2145 | GLU1775 | 0.65 | 16 | LEU2184 | LEU1817 | 0.54 | 22 | SER2191 | LEU1824 | 0.54 |
| 5 | GLY2136 | LYS1767 | 0.54 | 11 | ASP2146 | LEU1776 | 0.66 | 17 | ARG2185 | ARG1818 | 0.63 | 23 | LYS2193 | ARG1826 | 0.55 |
| 6 | ILE2137 | MET1768 | 0.64 | 12 | SER2147 | ASP1777 | 0.66 | 18 | LYS2186 | SER1819 | 0.72 | | | | |

### Table S5: hFAS-I KR and MtbFAS-I KR mapped residues table

| Sr. | hFAS-I | MtbFAS-I | Qh | Sr. | hFAS-I | MtbFAS-I | Qh | Sr. | hFAS-I | MtbFAS-I | Qh | Sr. | hFAS-I | MtbFAS-I | Qh |
|---|---|---|---|---|---|---|---|---|---|---|---|---|---|---|---|
| 1 | PHE1896 | ILE2108 | 0.63 | 33 | SER1942 | VAL2155 | 0.61 | 65 | GLN1989 | GLU2216 | 0.65 | 97 | GLY2052 | ALA2287 | 0.5 |
| 2 | GLY1897 | ALA2109 | 0.64 | 34 | THR1943 | ALA2156 | 0.6 | 66 | ASP1990 | MET2217 | 0.65 | 98 | PRO2054 | SER2290 | 0.66 |
| 3 | LEU1898 | ALA2110 | 0.62 | 35 | SER1944 | ALA2157 | 0.56 | 67 | VAL1991 | GLU2218 | 0.64 | 99 | GLY2055 | LEU2291 | 0.62 |
| 4 | GLU1899 | SER2111 | 0.63 | 36 | ASN1945 | ASN2158 | 0.5 | 68 | CYS1992 | MET2219 | 0.69 | 100 | LEU2056 | ALA2292 | 0.57 |
| 5 | LEU1900 | VAL2112 | 0.64 | 37 | SER1948 | SER2161 | 0.57 | 69 | LYS1993 | LYS2220 | 0.62 | 101 | ALA2057 | HIS2293 | 0.55 |
| 6 | ALA1901 | VAL2113 | 0.67 | 38 | LEU1949 | TYR2162 | 0.61 | 70 | PRO1994 | VAL2221 | 0.57 | 102 | VAL2058 | ALA2294 | 0.59 |
| 7 | GLN1902 | ALA2114 | 0.66 | 39 | GLU1950 | SER2163 | 0.64 | 71 | LYS1995 | LEU2222 | 0.67 | 103 | GLN2059 | LEU2295 | 0.61 |
| 8 | TRP1903 | ARG2115 | 0.65 | 40 | GLY1951 | ASP2164 | 0.67 | 72 | TYR1996 | LEU2223 | 0.73 | 104 | TRP2060 | ILE2296 | 0.61 |
| 9 | LEU1904 | LEU2116 | 0.64 | 41 | ALA1952 | VAL2165 | 0.71 | 73 | SER1997 | TRP2224 | 0.74 | 105 | GLY2061 | GLY2297 | 0.56 |
| 10 | ILE1905 | LEU2117 | 0.65 | 42 | ARG1953 | ASP2166 | 0.66 | 74 | GLY1998 | ALA2225 | 0.66 | 106 | ALA2062 | TRP2298 | 0.64 |
| 11 | GLN1906 | ASP2118 | 0.64 | 43 | GLY1954 | ALA2167 | 0.64 | 75 | THR1999 | VAL2226 | 0.67 | 107 | ILE2063 | THR2299 | 0.62 |
| 12 | ARG1907 | GLY2119 | 0.6 | 44 | LEU1955 | LEU2168 | 0.62 | 76 | LEU2000 | GLN2227 | 0.65 | 108 | GLY2064 | ARG2300 | 0.61 |
| 13 | GLY1908 | GLY2120 | 0.63 | 45 | ILE1956 | VAL2169 | 0.65 | 77 | ASN2001 | ARG2228 | 0.73 | 109 | ASP2065 | GLY2301 | 0.5 |
| 14 | VAL1909 | ALA2121 | 0.65 | 46 | ALA1957 | GLU2170 | 0.58 | 78 | LEU2002 | LEU2229 | 0.72 | 110 | THR2072 | MET2305 | 0.51 |
| 15 | LYS1911 | THR2122 | 0.57 | 47 | GLU1958 | TRP2171 | 0.52 | 79 | ASP2003 | ILE2230 | 0.5 | 111 | ASN2076 | VAL2312 | 0.5 |
| 16 | LEU1912 | VAL2123 | 0.62 | 48 | ALA1959 | ILE2172 | 0.54 | 80 | ARG2004 | GLY2231 | 0.51 | 112 | ILE2079 | VAL2315 | 0.64 |
| 17 | VAL1913 | ILE2124 | 0.68 | 49 | ALA1960 | GLY2173 | 0.54 | 81 | VAL2005 | GLY2232 | 0.65 | 113 | PRO2085 | THR2322 | 0.58 |
| 18 | LEU1914 | ALA2125 | 0.7 | 50 | GLY1963 | GLN2191 | 0.6 | 82 | THR2006 | LEU2233 | 0.51 | 114 | GLN2086 | TYR2323 | 0.6 |
| 19 | THR1915 | THR2126 | 0.69 | 51 | PRO1964 | THR2192 | 0.6 | 83 | ALA2009 | ILE2236 | 0.56 | 115 | ARG2087 | SER2324 | 0.56 |
| 20 | SER1916 | THR2127 | 0.64 | 52 | VAL1965 | PRO2193 | 0.6 | 84 | CYS2010 | GLY2237 | 0.57 | 116 | MET2088 | THR2325 | 0.53 |
| 21 | ARG1917 | SER2128 | 0.6 | 53 | GLY1966 | THR2194 | 0.57 | 85 | LEU2013 | LEU2246 | 0.55 | 117 | CYS2091 | MET2328 | 0.58 |
| 22 | SER1918 | LYS2129 | 0.6 | 54 | GLY1967 | LEU2195 | 0.59 | 86 | PHE2016 | VAL2248 | 0.62 | 118 | VAL2094 | LEU2331 | 0.51 |
| 23 | ALA1926 | LEU2135 | 0.58 | 55 | VAL1968 | LEU2196 | 0.63 | 87 | VAL2017 | VAL2249 | 0.54 | 119 | LEU2095 | LEU2332 | 0.52 |
| 24 | LYS1927 | ALA2136 | 0.53 | 56 | PHE1969 | PHE2197 | 0.65 | 88 | VAL2018 | LEU2250 | 0.56 | 120 | LEU2099 | CYS2336 | 0.5 |
| 25 | VAL1929 | TYR2138 | 0.55 | 57 | ASN1970 | PRO2198 | 0.69 | 89 | PHE2019 | PRO2251 | 0.59 | 121 | VAL2105 | LYS2350 | 0.5 |
| 26 | ARG1930 | ARG2139 | 0.52 | 58 | LEU1971 | PHE2199 | 0.65 | 90 | SER2020 | GLY2252 | 0.64 | 122 | LEU2106 | ALA2351 | 0.55 |
| 27 | ARG1931 | THR2140 | 0.53 | 59 | ALA1972 | ALA2200 | 0.65 | 91 | ARG2026 | ASP2262 | 0.53 | 123 | SER2107 | ASP2352 | 0.54 |
| 28 | VAL1937 | ALA2150 | 0.5 | 60 | VAL1973 | ALA2201 | 0.6 | 92 | GLY2027 | GLY2263 | 0.57 | 124 | SER2108 | LEU2353 | 0.57 |
| 29 | GLN1938 | ALA2151 | 0.57 | 61 | VAL1974 | PRO2202 | 0.57 | 93 | PHE2036 | SER2270 | 0.51 | 125 | PHE2109 | THR2354 | 0.54 |
| 30 | VAL1939 | LEU2152 | 0.63 | 62 | GLU1986 | SER2213 | 0.6 | 94 | ALA2037 | ALA2271 | 0.5 | 126 | VAL2110 | GLY2355 | 0.54 |
| 31 | GLN1940 | TRP2153 | 0.66 | 63 | PHE1987 | ARG2214 | 0.53 | 95 | SER2039 | ASP2273 | 0.51 | | | | |
| 32 | VAL1941 | LEU2154 | 0.66 | 64 | PHE1988 | ALA2215 | 0.62 | 96 | ALA2040 | ALA2274 | 0.61 | | | | |

### Table S6: hFAS-I DH and MtbFAS-I DH mapped residues table

| S.N | hFAS-I | MtbFAS-I | QH | S.N | hFAS-I | MtbFAS-I | QH | S.N | hFAS-I | MtbFAS-I | QH | S.N | hFAS-I | MtbFAS-I | QH |
|---|---|---|---|---|---|---|---|---|---|---|---|---|---|---|---|
| 1 | ALA888 | PRO1063 | 0.51 | 17 | VAL954 | ALA1149 | 0.56 | 33 | SER1037 | VAL1242 | 0.57 | 49 | ARG1085 | VAL1288 | 0.54 |
| 2 | VAL913 | ASP1100 | 0.52 | 18 | GLN960 | PHE1155 | 0.52 | 34 | ILE1038 | THR1243 | 0.52 | 50 | VAL1086 | ASP1289 | 0.62 |
| 3 | LEU919 | VAL1105 | 0.6 | 19 | GLY1015 | ARG1189 | 0.55 | 35 | PRO1049 | LEU1255 | 0.54 | 51 | THR1087 | VAL1290 | 0.63 |
| 4 | HIS920 | VAL1106 | 0.61 | 20 | SER1017 | ASP1190 | 0.62 | 36 | THR1050 | VAL1256 | 0.57 | 52 | VAL1088 | ALA1291 | 0.62 |
| 5 | SER932 | THR1117 | 0.5 | 21 | GLY1018 | VAL1191 | 0.5 | 37 | ARG1051 | GLY1257 | 0.52 | 53 | ALA1089 | ALA1292 | 0.59 |
| 6 | LEU933 | VAL1118 | 0.56 | 22 | ARG1019 | THR1192 | 0.57 | 38 | ILE1055 | ALA1260 | 0.5 | 54 | GLY1091 | VAL1299 | 0.54 |
| 7 | GLU934 | THR1119 | 0.57 | 23 | LEU1020 | ILE1193 | 0.56 | 39 | HIS1056 | ARG1261 | 0.55 | 55 | VAL1092 | MET1300 | 0.55 |
| 8 | VAL935 | ALA1120 | 0.56 | 24 | LEU1021 | THR1194 | 0.51 | 40 | ILE1057 | PHE1262 | 0.51 | 56 | HIS1093 | SER1301 | 0.58 |
| 9 | ARG936 | THR1121 | 0.55 | 25 | TRP1022 | ALA1195 | 0.5 | 41 | VAL1075 | GLU1271 | 0.53 | 57 | ILE1094 | ALA1302 | 0.61 |
| 10 | GLU945 | VAL1139 | 0.53 | 26 | MET1030 | SER1235 | 0.53 | 42 | ALA1076 | VAL1272 | 0.53 | 58 | SER1095 | SER1303 | 0.57 |
| 11 | VAL946 | VAL1140 | 0.52 | 27 | ASP1031 | ALA1236 | 0.57 | 43 | ASP1077 | ASP1273 | 0.54 | 59 | LEU1097 | ALA1304 | 0.63 |
| 12 | SER947 | THR1141 | 0.53 | 28 | THR1032 | ALA1237 | 0.56 | 44 | VAL1078 | PHE1274 | 0.56 | 60 | HIS1098 | ARG1305 | 0.62 |
| 13 | GLY950 | GLY1145 | 0.56 | 29 | MET1033 | ALA1238 | 0.58 | 45 | VAL1079 | ARG1275 | 0.6 | 61 | THR1099 | LEU1306 | 0.57 |
| 14 | ASN951 | ALA1146 | 0.57 | 30 | LEU1034 | GLN1239 | 0.59 | 46 | VAL1080 | VAL1276 | 0.62 | 62 | GLU1100 | ALA1307 | 0.61 |
| 15 | LEU952 | VAL1147 | 0.6 | 31 | GLN1035 | HIS1240 | 0.56 | 47 | SER1081 | GLU1277 | 0.62 | | | | |
| 16 | VAL953 | ILE1148 | 0.56 | 32 | MET1036 | ALA1241 | 0.56 | 48 | ARG1082 | ARG1278 | 0.59 | | | | |
